# Multi-phosphorylation reaction and clustering tune Pom1 gradient mid-cell levels according to cell size

**DOI:** 10.1101/546424

**Authors:** Veneta Gerganova, Charlotte Floderer, Anna Archetti, Laetitia Michon, Lina Carlini, Thaïs Reichler, Suliana Manley, Sophie G Martin

## Abstract

Protein concentration gradients convey information at a distance from the source to both pattern developing organisms and organize single cells. In the rod-shaped cells of *Schizosaccharomyces pombe*, the DYRK-family kinase Pom1 forms concentration gradients with maxima at the cell poles. Pom1 controls the timing of mitotic entry by inhibiting the SAD-family kinase Cdr2, which forms stable membrane-associated nodes at mid-cell. Pom1 gradients rely on membrane association regulated by a phosphorylation-dephosphorylation cycle and lateral diffusion modulated by clustering. Whether the graded pattern directly alters Pom1 medial levels has been controversial. Here, using a combination of quantitative imaging approaches, including single particle tracking PALM and TIRF microscopy, we find that individual Pom1 molecules do not bind the membrane long enough to diffuse from cell pole to cell middle. Instead we propose they exchange within longer-lived clusters that form the functional gradient units. By creating an allelic series progressively blocking auto-phosphorylation, we show that multi-phosphorylation shapes and buffers the gradient to control the cortical mid-cell Pom1 levels, which represent the critical pool regulating Cdr2. Specific imaging of this cortical pool by TIRF microscopy demonstrates that more Pom1 overlaps with Cdr2 nodes in short than long cells, consistent with Pom1 inhibition of Cdr2 decreasing with cell growth. We conclude that Pom1 gradients modulate Pom1 mid-cell levels according to cell size.

## Introduction

In many organisms and cell types, graded protein patterns provide positional information. This is true from the smallest bacteria, where polar gradients of protein activity define the position of the division apparatus (Kretschmer and Schwille, 2016), to the largest multicellular organisms, where morphogen concentration gradients define regions of gene expression during development (Briscoe and Small, 2015). Although mechanisms of gradient formation vary, in all systems the graded patterns are thought to convey information at a distance from the source.

In fission yeast, concentration gradients formed by the DYRK-family kinase Pom1 have received considerable attention, due to the role of Pom1 in regulating the timing of mitotic entry and thus cell size at division (Martin and Berthelot-Grosjean, 2009; Moseley et al., 2009). Pom1 gradients are nucleated at cell poles upon dephosphorylation by a type I phosphatase complex whose regulatory subunit Tea4 is delivered by microtubules (Hachet et al., 2011; Martin et al., 2005; Tatebe et al., 2005). Dephosphorylation of Pom1 reveals a lipid-binding activity that maps to a 200-aa region in its N-terminal half. At the plasma membrane, Pom1 forms small clusters and is thought to laterally diffuse (Hachet et al., 2011; Saunders et al., 2012). It also undergoes autophosphorylation reactions that promote its detachment from the membrane, leading to the graded pattern (Hachet et al., 2011).

An interesting feature of Pom1 gradients is a noise correction mechanism that compensates for large variations in protein concentration at the cell poles, which can vary up to 4-fold (Hersch et al., 2015; Saunders et al., 2012). In a simple diffusive gradient, the decay length (the distance at which the concentration is reduced to a certain fraction of its amplitude) is independent of the amplitude at the pole. In contrast, the Pom1 gradient is corrected by varying the slope of the gradient decay: gradients with higher Pom1 concentration have a steeper decay, while those with lower Pom1 concentration have a flatter decay. Two models have been proposed to explain the source of this correction. One model suggests gradient buffering is the consequence of concentration-dependent inter-molecular phosphorylation, which promotes Pom1 detachment from the membrane (Hersch et al., 2015). This model also explains the correction for the even larger variations in levels of Tea4 concentration at cell poles. A second model hypothesizes that buffering results from concentration-dependent clustering of Pom1, with higher concentrations leading to larger, slower diffusive clusters, causing a traffic-jam effect at the cell tips (Saunders et al., 2012). However, direct experimental evidence testing these models is scarce.

Pom1 has two physiological functions. First, Pom1 provides spatial information for division site positioning: *pom1Δ* cells divide off-centre (Bähler and Nurse, 2001; Bähler and Pringle, 1998). Second, Pom1 provides temporal information to regulate the timing of mitotic entry: *pom1Δ* cells divide precociously at an aberrantly small size (Martin and Berthelot-Grosjean, 2009; Moseley et al., 2009). For both functions, Pom1 directly phosphorylates the SAD-family kinase Cdr2, but on different residues (Bhatia et al., 2013; Rincon et al., 2014). Pom1 function in division site placement likely also involves additional substrates. To delay mitotic commitment, Pom1 phosphorylates the C-terminus of Cdr2 (Bhatia et al., 2013; Deng et al., 2014), which antagonizes the activating phosphorylation of the Cdr2 kinase domain by the cytosolic CaMKK Ssp1 (Deng et al., 2014). Cdr2 localizes at the mid-cell cortex, where it forms large, stable clusters called nodes, which contain many other proteins including a second SAD-family kinase Cdr1 (Martin and Berthelot-Grosjean, 2009; Morrell et al., 2004; Moseley et al., 2009). The signal relay between Cdr2 and Cdr1 is not yet elucidated, but the output is an inhibitory phosphorylation of Wee1 kinase, which itself exerts direct inhibitory activity on the sole cyclin-dependent kinase CDK1 (Kanoh and Russell, 1998; Young and Fantes, 1987). In contrast to the stable Cdr1 and Cdr2 association to the nodes, Wee1 visits are only transient (Allard et al., 2018; Moseley et al., 2009). These visits increase in frequency and duration as cells grow, consistent with the idea that Wee1 is inactivated in longer cells to permit CDK1 activation and mitotic entry.

Although genetic and biochemical evidence have firmly established Cdr2 as Pom1 substrate, there has been much debate on where the Pom1-Cdr2 interaction takes place, and whether the strength of this interaction varies in the course of a single cell cycle. Initial work proposed Pom1 gradients as a means to measure cell size, because total fluorescence measurements of Pom1-GFP along cell length revealed higher medial fluorescence in short than long cells (Martin and Berthelot-Grosjean, 2009; Moseley et al., 2009). This led to the model that Pom1 inhibits Cdr2 in short cells, but that inhibition is relieved upon attaining sufficient cell size, thus coupling cell size with mitotic entry. Two lines of evidence indicate that Pom1 activity on Cdr2 indeed varies with cell size. First, the levels of Cdr2 phosphorylation by Ssp1 increase during G2, consistent with a progressive decrease in Pom1-dependent inhibition (Deng et al., 2014). Second, the frequency and duration of Wee1 visits to Cdr2 nodes increase as cells grow, with direct evidence showing that Pom1 suppresses Wee1 visits in short cells (Allard et al., 2018). In addition, Pom1 re-localization to cell sides, which is prominent upon glucose starvation, leads to strong mitotic delay (Kelkar and Martin, 2015). However, subsequent analyses of cortical fluorescence profiles on confocal mid-plane images failed to detect significant differences in the levels of Pom1-GFP at mid-cell between short and long cells, raising questions about the previously proposed model (Bhatia et al., 2013; Pan et al., 2014). The apparently invariant Pom1 mid-cell levels in cells of various lengths led to the suggestion that Pom1 may control Cdr2 activity elsewhere (Bhatia et al., 2013), or is not involved in cell size homeostasis (Pan et al., 2014), in agreement with the observation that *pom1Δ* cells retain homeostatic capacity (Wood and Nurse, 2013). Because the number of Cdr2 nodes at mid-cell increases with cell surface growth, this also led to the suggestion that Cdr2, not Pom1, may be the critical cell size sensor in the pathway (Pan et al., 2014). Thus, there is currently a controversy between the invariant Pom1 levels at mid-cell and the size-dependent effect of Pom1 on Cdr2 function.

In this work, we have coupled generation of a systematic *pom1* mutant allelic series with a wide range of imaging methods – including single molecule super-resolution PALM imaging and tracking, confocal and TIRF microscopy – to obtain quantitative information on the patterning of Pom1 gradients. This enabled three major findings: first, Pom1 gradients are primarily shaped by phosphorylation-mediated detachment with clusters acting as the relevant membrane-associated unit; second, Pom1 regulates Cdr2 for mitotic entry at the mid-cell cortex; third, TIRF imaging reveals significantly higher levels of Pom1 at the mid-cell cortex in short cells.

## Results

### Membrane diffusion and dissociation of Pom1

We investigated the diffusion and membrane dissociation dynamics of single molecules of Pom1 in *S*. *pombe* cells. Cells were prepared either on a flat agarose pad and imaged horizontally along their long axis, or on a micropatterned surface imprinted with holes, where cells oriented vertically for cell pole imaging (Figure 1A, Figure 1 – supplement 1). We relied on photoconversion, localization, and tracking of single fluorescent proteins in living cells, in our case the Pom1-mEos3.2 fusion expressed from the native genomic locus. Single molecules of Pom1-mEos3.2 were observed along the side or pole of the cell, with a higher density of localizations at the poles consistent with the known Pom1 density gradient, and tracked by monitoring their position over time to produce single molecule trajectories (Figure 1B) (Manley et al., 2008).

**Figure 1.**
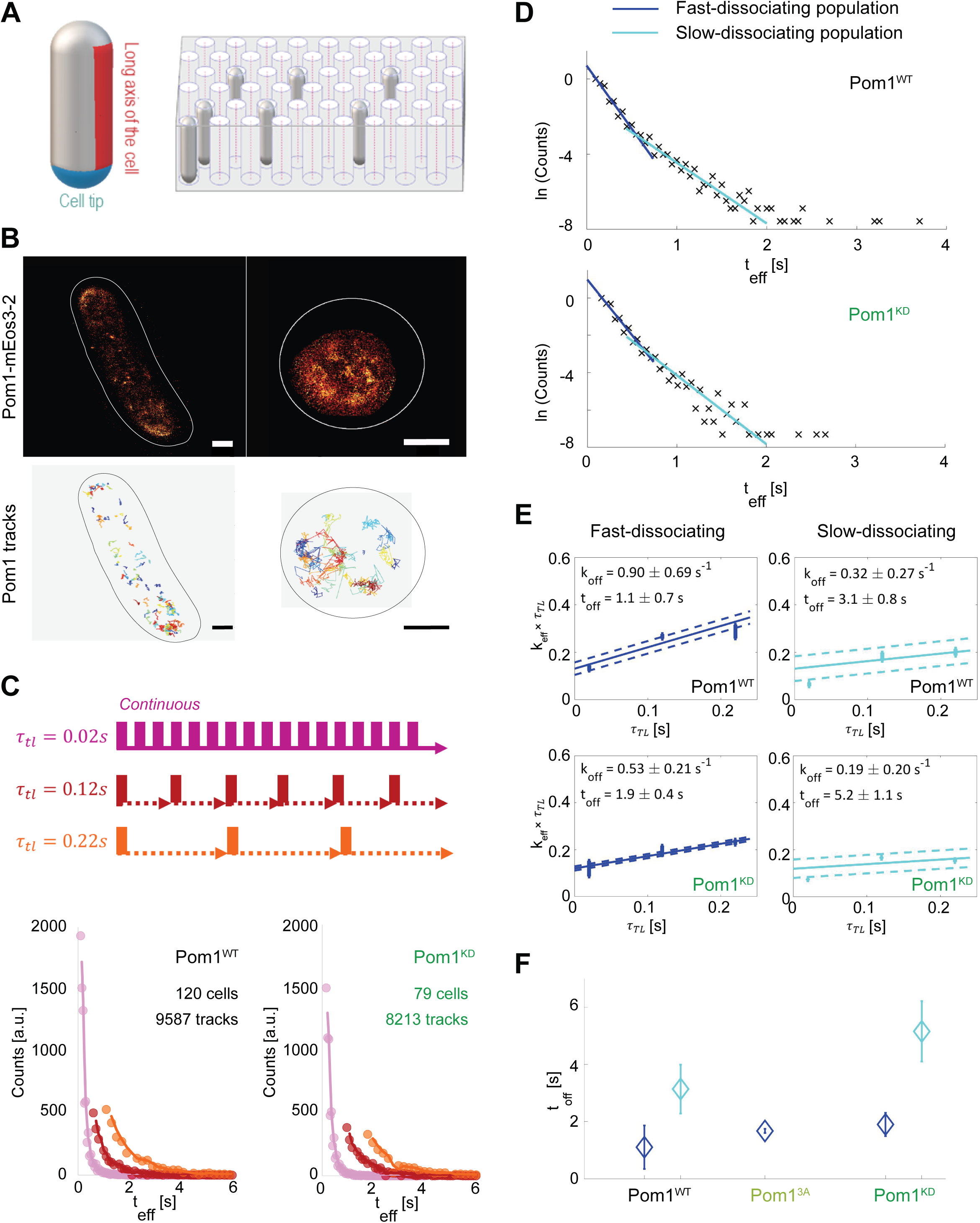
Pom1 dissociation dynamics. **A.** (left) Scheme of the cell regions imaged on flat pads (cell side, red) or vertical molds (cell pole, blue). (right) Scheme of the mold used for vertical immobilization in agar. **B.** PALM reconstruction of Pom1 on the cell side (top left) or the cell pole (top right). White lines correspond to cell boundaries. Corresponding sptPALM tracks on the side (bottom left) and on the pole (bottom right). Scale bar 1 µm. **C.** Scheme for time-lapse imaging experiments. Every frame is recorded with a 20ms exposure time (solid rectangles). A time delay (dashed arrows) is introduced between each pair of consecutive frames. The time-lapse period (τ_*TL*_) is the sum of the integration time and the time delay. Effective residence time distributions for Pom1^WT^ and Pom1^KD^, colour correspond to different τ_*TL*_. **D.** Residence time distributions for Pom1^WT^ and Pom1^KD^, fit with a bi-exponential decay: fast (blue) and slow (cyan). **E.** Effective rate constant as a function of time lag condition (symbols) for fast (blue) and slow (cyan) populations. Solid lines correspond to a weighted linear fit according to Eq.2. Error bars correspond to the weights associated to each data point (S.D. from the fit of the exponential distribution of the residence time obtained according to Eq. 1). **F.** Comparison of the Pom1 residence time obtained for each subpopulation (fast in blue and slow in cyan) for Pom1^WT^, Pom1^3A^ and Pom1^KD^ strains. Error bar corresponds to the standard deviation of the parameter extracted from the weighted linear fit of panel 1E.

To determine the dissociation rate of single molecules, we performed time-lapse imaging on horizontally oriented cells with several lag times *τ*_TL_ (Gebhardt et al., 2013) (Figure 1C). This allows the effects of dissociation and photobleaching to be analytically separated, since varying the lag time will vary the contribution of photobleaching and therefore the effective residence time (t_eff_) (Figure 1C), while the actual residence time of molecules remains unchanged. Interestingly, we observed that the residence time exhibits a multimodal distribution, which could be fitted with a bi-exponential decay corresponding to short and long residence times, or fast and slow dissociation rates (Figure 1D, Figure 1 – supplement 2A). The majority of Pom1 molecules (76%) comprise the fast-dissociating part, while the remaining (23%) dissociate more slowly. This last population contained few molecules, but its long tail could be further explained by the presence of two or more slowly dissociating sub-populations.

To understand the contribution of Pom1 auto-phosphorylation in membrane detachment, we analysed the time-lapse data of Pom1 binding using both Pom1^WT^-mEos3.2 and kinase-dead Pom1^KD^-mEos3.2 to extract binding times (t_off_) and dissociation rates (k_off_ =1/t_off_) (Figure 1E). Both Pom1^WT^ and Pom1^KD^ showed fast and slow-dissociating populations, with a similar fold difference in dissociation rates between the two populations (2.8x and 2.7x, respectively). Interestingly, when we compared the fast-dissociating populations in the two strains, the Pom1^WT^ population dissociated 1.7x faster than the Pom1^KD^ population 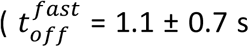 and 1.9 ± 0.4 s for Pom1^WT^ and Pom1^KD^, respectively). The slowly dissociating population showed a similar trend (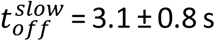 and 5.2 ± 1.1 s for Pom1^WT^ and Pom1^KD^, respectively) (Figure 1E-F). We note that an even slower sub-population may exist, shown as the tail distribution in Fig. 1D, but this represents a very small fraction of Pom1 molecules (<1%) for which we lack sufficient number of tracks to extract a reliable dissociation rate.

Thus, Pom1 activity, which leads to auto-phosphorylation, promotes a faster dissociation rate of Pom1 from the membrane, in agreement with previous biochemical observations (Hachet et al., 2011). Furthermore, both Pom1^WT^ and Pom1^KD^ dissociation kinetics exhibit at least two distinct populations perhaps corresponding to different multimerization states.

### Localization and diffusion of Pom1

To study the lateral diffusion of Pom1 at the plasma membrane, single fluorescent Pom1 proteins were tracked (Fig. 1B) and analyzed to extract their diffusion coefficients. We found that the track duration t_eff_ and diffusion coefficient D_eff_ were inversely correlated (Fig. 2A), with shorter tracks exhibiting faster diffusion and longer tracks exhibiting slower diffusion. Since long residence times imply a slower dissociation time, slowly diffusing Pom1 molecules present slow dissociation dynamics, and vice versa. This suggests that in larger, slowly diffusing clusters, Pom1 remains more stably associated with the membrane.

**Figure 2.**
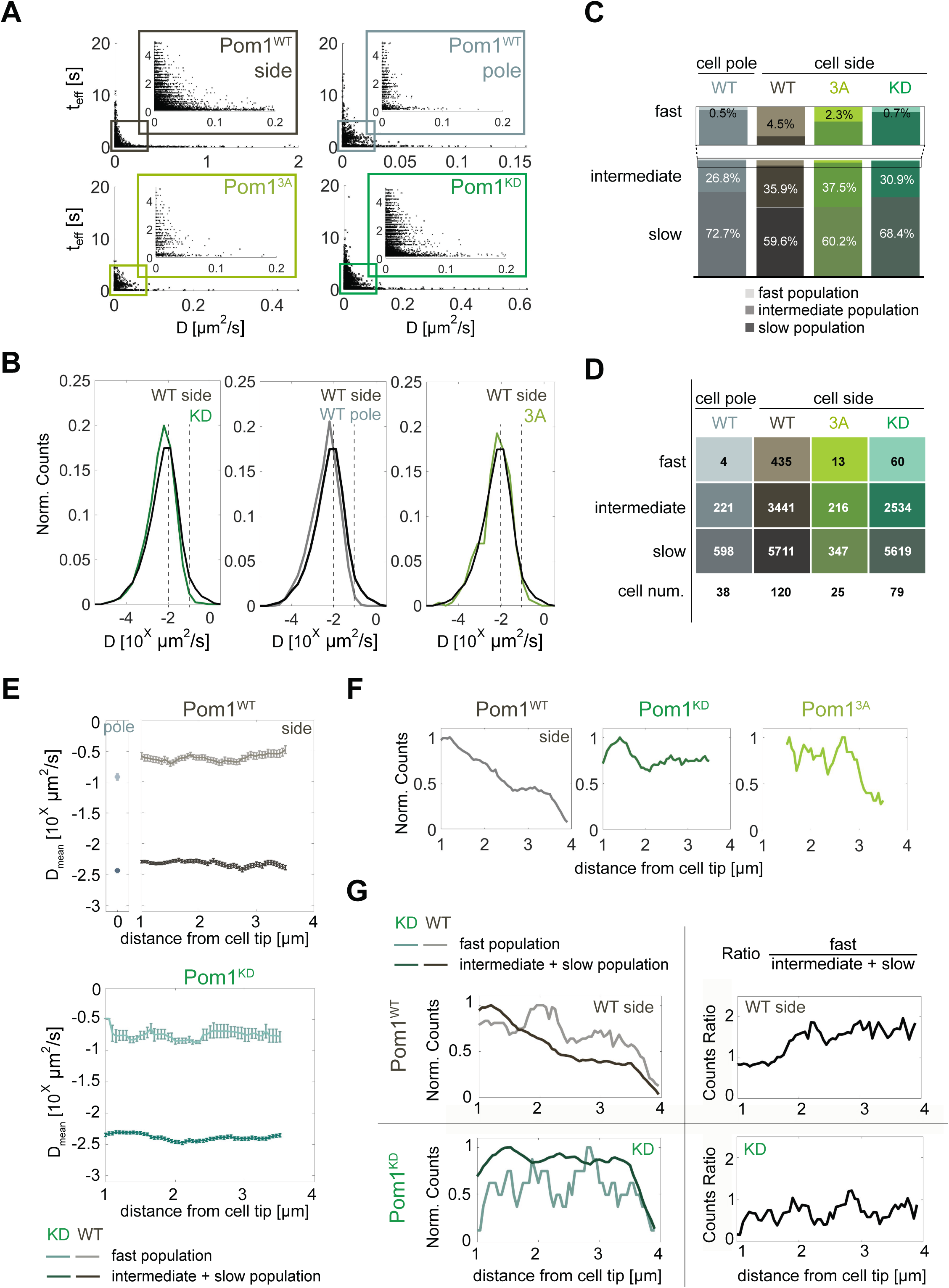
Pom1 diffusion dynamics. **A.** Track length as a function of diffusion coefficients of the tracks for Pom1^WT^ (WT) at cell sides (dark grey) and cell poles (light grey), Pom1^3A^ (3A; light green), Pom1^KD^ (KD; green-cyan). Pom1^3A^ and Pom1^KD^ were imaged at cell sides. **B.** Distribution of diffusion coefficients of all Pom1 molecules tracked in Pom1^WT^, Pom1^3A^ and Pom1^KD^ at cell sides, and Pom1^WT^ at cell poles. Thresholds used in panels C-G are shown by the dashed lines. **C.** Proportion of fast (D ≥ 10^−1^ µm^2^/s; light), intermediate (10^−2^ ≤ D < 10^−1^ µm^2^/s; medium color) and slow (D < 10^−2^ µm^2^/s; dark) populations for Pom1^WT^, Pom1^3A^ and Pom1^KD^ at cell sides, and Pom1^WT^ at cell poles. **D.** Number of tracks and cells for each condition. **E.** Average diffusion coefficient as a function of the distance from the pole for fast (D ≥ 0.1µm^2^/s; light color) and combined intermediate and slow (dark) populations for Pom1^WT^ (D < 0.1µm^2^/s; top panels) and Pom1^KD^ (bottom) strains. Error bar corresponds to standard error of the mean. **F.** Evolution of count of all tracks along the cell length normalized by the maximum occurrence for Pom1^WT^, Pom1^KD^ and Pom1^3A^. **G.** Evolution of the number of tracks for fast (D ≥ 0.1µm^2^/s; light color) and combined intermediate and slow (D < 0.1µm^2^/s; dark) populations along the cell length for Pom1^WT^ and Pom1^KD^ strains (left panels) and their ratio (right panel) for Pom1^KD^ and Pom1^KD^ strains. The color coding dark grey = Pom1^WT^ at cell sides, light grey = Pom1^WT^ at cell poles, green-cyan = Pom1^KD^, and light green = Pom1^3A^ is used throughout the figure.

Pom1^WT^ and Pom1^KD^ exhibit a broad distribution of diffusion coefficients (Fig. 2B), with the distribution of Pom1^WT^ at cell sides shifted towards slightly faster diffusion compared with Pom1^KD^. To better understand this difference, we defined two thresholds at D ≥ 10^−1^ µm^2^/s and D ≥ 10^−2^ µm^2^/s, which separate molecules into three sub-populations of fast (*D* ≥ 10^−1^ µm^2^/s), intermediate (10^−2^ ≤ *D* < 10^−1^ µm^2^/s) and slow-diffusing molecules (*D* < 10^−2^ µm^2^/s) (Figure 2B). Note that in this analysis, all fast-diffusing molecules are also fast-dissociating, but intermediate and slow-diffusing molecules can exhibit either fast- or slow-dissociating behaviours. Interestingly, there was a substantially higher proportion of fast-diffusing molecules for Pom1^WT^ than Pom1^KD^, which also on average diffused faster (*D*_*mean*_ = 0.31 ± 0.01 µm^2^/s for Pom1^WT^ and 0.21 ± 0.02 µm^2^/s Pom1^KD^; Figure 2C-E). We note that the average diffusion rate of these three populations did not vary along cell length with distance from cell poles (Fig 2E). These analyses were robust to changes in threshold choice (Figure 2 – supplement 1). Thus, the fast population of Pom1^WT^ diffuses faster than Pom1^KD^.

We then performed the same analysis on Pom1^WT^ at the cell pole using cells oriented vertically (Fig. 1A). Interestingly, the distribution of diffusion coefficients was shifted towards slower values compared to Pom1^WT^ at the cell sides (Fig. 2B). Indeed, when using the same thresholds as above, a much smaller proportion of molecules were fast-diffusing compared with the sides (Fig. 2C; Figure 2 – supplement 1A-C), very similar to the proportion observed for Pom1^KD^. This is consistent with Pom1 being mainly in its dephosphorylated state at the cell pole, in agreement with the presence of the Tea4-PP1 phosphatase at this location (Hachet et al., 2011).

We then tracked the evolution of the number of events along cell length starting from the cell tip. The overall number of events decreased for Pom1^WT^ but not Pom1^KD^, consistent with their described localization patterns (Figure 2F). Interestingly, when we considered fast, intermediate and slow populations separately, we found that Pom1^WT^ exhibits a change in the relative proportion of molecules: the proportion of fast diffusive Pom1 remained relatively constant all along the gradient, but the proportion of intermediate and slow diffusive Pom1 decreased, resulting in a relative increase in the fast population (Figure 2G, Figure 2 – supplement 1E-G). In contrast, the proportions of fast, intermediate and slow diffusive populations of Pom1^KD^ remain balanced all along the gradient, leading to a nearly constant ratio.

In summary, these measurements provide two important insights: First, the measured binding times and diffusion rates indicate that individual Pom1 molecules cover on average a small distance before detaching from the membrane. Let’s consider the fast-diffusing molecules. These represent 4.5% of the population, diffuse on average at 0.31µm^2^/s and are all part of the 76% of molecules binding the membrane for an average time of 1.1 s. In fact, given the inverse correlation observed between diffusion and binding times, they likely bind the membrane for an even shorter time. From these values, we can estimate a maximum travelled distance of 0.8µm. Slower-diffusing molecules travel an even shorter distance before detaching. Thus, it is unlikely that individual molecules continuously track from cell pole to cell sides. Second, the shorter binding time of fast-diffusive molecules and their progressive increase in proportion at a distance from the cell pole for Pom1^WT^ but not Pom1^KD^ suggests the possibility this may be caused by progressive phosphorylation-dependent Pom1 detachment from larger clusters. This led us to examine more closely the mode of Pom1 attachment to the membrane and the role auto-phosphorylation plays to shape the gradient.

### Pom1 binds the plasma membrane through two distinct motifs

Previous work has shown that Pom1 interacts with lipids (Hachet et al., 2011) and a fragment containing amino acids 305 to 510 can efficiently bind the plasma membrane (Figure 3A, fragment #1). A BH-search prediction (Brzeska et al., 2010) performed for this fragment identified two potential membrane binding sites (Figure 3B). These two regions map to the most conserved sequences in the 305-510 fragment: the first (aa 437-444) is within a 22aa-long sequence (MB1; aa 423-444) that is identical in the four *Schizosaccharomyces* species (*S*. *pombe, S*. *octosporus, S*. *japonicus, and S*. *cryophilus*), the second (aa 480-492) falls within a weak amphipathic helix prediction (MB2; aa 477-494) (Figure 3A-C). To test the validity of these predictions, we constructed a series of shorter and/or mutagenized GFP-tagged Pom1 fragments integrated as single copy in wildtype and *pom1Δ* cells. The localization of all fragments tested was identical in wildtype and *pom1Δ* cells (Figure 3D and Figure 3- supplement 1).

**Figure 3.**
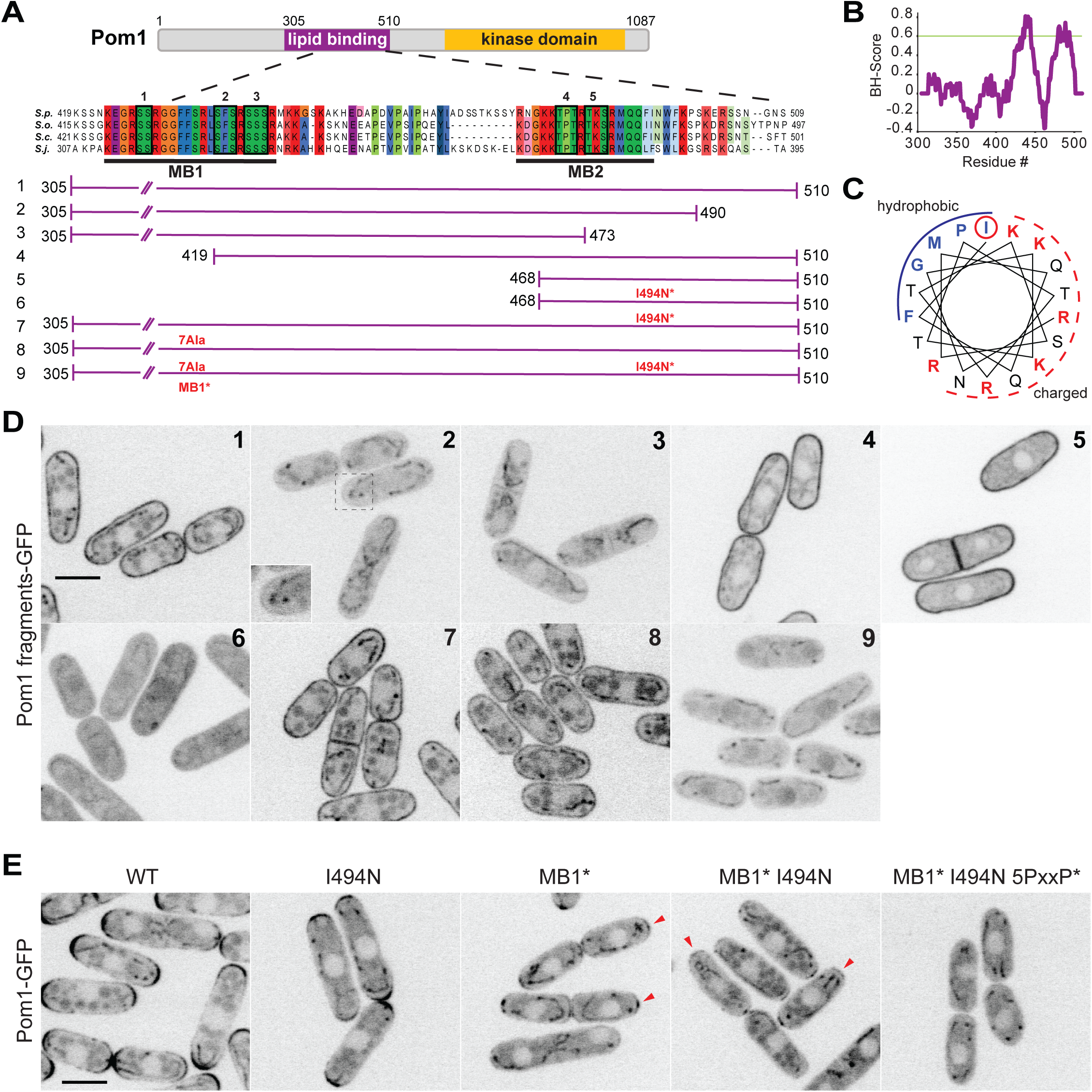
Two distinct regions define Pom1 membrane binding. **A.** Schematic representation of Pom1 with sequence homology alignment of the lipid-binding region shown for *S*. *pombe, S*. *octosporus, S*. *cryophilus,* and *S*. *japonicus*. Phosphorylation sites are indicated by black boxes and numbered. The conserved membrane binding region MB1 (aa423-444) and the amphipathic helix membrane binding region MB2 (aa477-494) are underlined. A schematic representation of the fragments used in the truncation analysis presented in D is shown below. **B.** BH-search prediction performed on Pom1 lipid-binding region (aa305-510) shows two peaks with a BH-score above 0.6, corresponding to membrane binding regions MB1 and MB2. **C.** Predicted amphipathic helix for MB2 with marked residue I494, targeted for mutagenesis I494N. **D.** Localization of GFP-tagged fragments 1 to 9 presented in A in wild type cells. Inset shows residual membrane localization for fragment #2. Scale bar 5μm. **E.** Localization of full-length Pom1^WT^-GFP and indicated mutants expressed at the native locus. Mutagenesis of the two membrane-binding regions and the 5 PxxP sites that mediate direct binding to Tea4 (Hachet et al., 2011) renders Pom1 cytosolic. Red arrowheads indicate residual cell tip localization in Pom1^MB1*^ and Pom1^MB1*-I494N^ mutants. Scale bar 5μm.

The Pom1 fragment spanning aa 305-510 localizes uniformly at the plasma membrane (fragment #1). Truncation of the C-terminal 20 aa (fragment 305-490aa, fragment #2), which cuts into the predicted amphipathic helix, resulted in a strong reduction (though not complete loss; see below) of Pom1 cortical localization. Progressive N-terminal truncations showed that a minimal 468-510aa fragment (fragment #5) containing the putative amphipathic helix was sufficient for membrane binding. In this fragment, converting the hydrophobic Ile494 to the polar, uncharged Asp residue disrupted membrane binding (I494N, fragment #6), indicating that Pom1 binds the membrane through the predicted amphipathic helix. However, the same point mutation in the full 305-510aa fragment (fragment #7) did not disrupt cortical localization, indicating the presence of a second lipid-binding domain. We then mutagenized 7 amino acids within the conserved MB1 region to alanine (generating the mutant allele MB1*, fragment #8), which also on its own did not perturb membrane binding. However, combining both the I494N and MB1* mutations within fragment 305-510 yielded a fully cytosolic localization (fragment #9). We conclude that Pom1 localization to the plasma membrane relies on two adjacent lipid-binding motifs.

To confirm the results from the fragment analysis, we introduced the same mutations in full-length Pom1 at the native genomic locus. Individual I494N and MB1* mutations led to a decrease of fluorescence intensity at the cell tip, which was exacerbated in the double mutant. Nevertheless, in this double mutant small Pom1 clusters were visible at the cell tips, likely due to Pom1 direct binding to the phosphatase regulatory subunit Tea4 through PxxP motifs (Hachet et al., 2011). Indeed, additional mutagenesis of the 5 previously identified PxxP motifs in combination with the MB1* and I494N mutations rendered Pom1 entirely cytosolic.

### A Pom1 allelic series shows additive features of multi-phosphorylation

Pom1 autophosphorylation promotes membrane detachment. From over 40 phosphorylation sites identified in silico and by mass-spectrometry analysis, combined mutation of 6 of these sites was previously shown to abolish the Pom1 gradient ((Hachet et al., 2011); note that each site contains up to 3 serines or threonines mutated in aggregate). Five of these sites are located in the 305-510aa fragment: number 1, 2, and 3 are within the conserved MB1 membrane-binding region, while 4 and 5 are located in the MB2 amphipathic helix (Figure 3A, sites indicated by black boxes). The 6^th^ one is distal to the kinase domain and was not directly investigated here.

To test the contribution of multi-phosphorylation for gradient shape and buffering, we generated a series of endogenously tagged phospho-blocking Pom1 alleles, carrying alanine substitution in 1, 2, 3 or 5 phosphorylation sites (Figure 4A, top row), and quantified Pom1 gradients by measuring the fluorescence profile at the cell cortex in medial plane confocal images. A decrease in the number of phosphorylation sites led to a gradual flattening of the gradient shape manifested in a decrease of Pom1 intensity at the cell tip and an increase at the cell middle (Figure 4B-C, left panels). The gradual change of gradient shape indicates the additive nature of multiple autophosphorylation events and reveals that no particular site contributes to Pom1 gradient shape more than another. The gradient shape of the Pom1^5A^ mutant was indistinguishable from those of the previously described non-phosphorylatable Pom1^6A^ and inactive Pom1^KD^ (Hachet et al., 2011), indicating that these five sites represent the principal sites modulating Pom1 localization.

**Figure 4.**
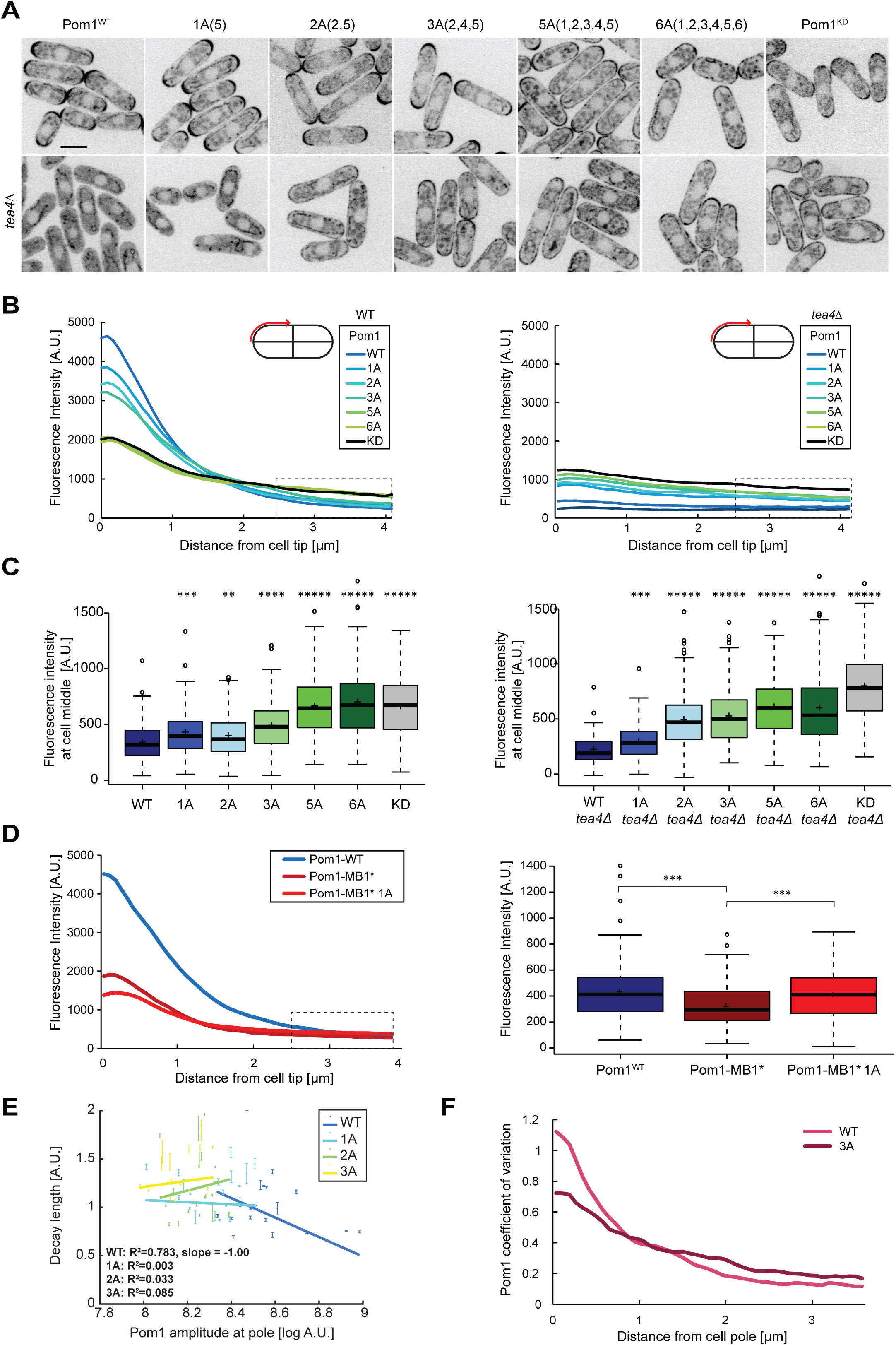
Pom1 gradient shape and robustness depend on multisite auto-phosphorylation. **A.** Medial plane confocal images of a series of Pom1 phospho-blocking alleles in otherwise wild type (top row) and *tea4Δ* cells (bottom row). Scale bar 5μm. **B.** Fluorescence intensity plots of cortical gradient profiles, collected from time-average medial plane confocal images as shown on the schematic. Left: wild type background; right: *tea4Δ* cells. Graphs show averages of 240 gradient profiles per strain, n=3 experiments, 20 cells per experiment. Individual experiments are shown in Figure 4 – supplement 1. The dashed box shows the region selected for Pom1 intensity measurements at mid-cell shown in panel C. **C.** Mean Pom1 fluorescence intensity levels at cell middle, extracted from the last 1.5µm of profiles shown in panel B. Left: wild type background, right: *tea4Δ* cells. **D.** Cortical gradient profiles of Pom1^WT^, Pom1^MB1*^ and Pom1^MB1*-1A(5)^ (left) and corresponding quantification of Pom1 intensity at cell middle (right), as in panels B and C. Graphs show averages of 160 gradient profiles per strain, n=2 experiments, 20 cells per experiment. **E.** Decay length plotted against Pom1 amplitude at the cell pole. Each dot represents an average gradient profile from bin sorting of 5%. Error bars correspond to the percentage of fitting quality for each dot. **F.** Coefficients of variation for Pom1^WT^ and Pom1^3A^ from the 240 gradient profiles shown in B. Means indicated by plus sign, error bars: SD, statistical significance measured against wild type unless otherwise indicated by t-test with unequal variances. **, p=0.0001, *** p=10^−8^, **** p≤10^−18^, ***** p≤10^−40^.

To further probe the influence of phosphorylation on Pom1 membrane binding, we used single molecule time-lapse imaging of Pom1^3A^-mEos3.2 to extract dissociation and diffusion rates. The fast-detaching population of Pom1^3A^ molecules showed membrane binding times intermediate between Pom1^WT^ and Pom1^KD^ (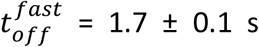 Figure 1F, Figure 1 – supplement 2B). The number of tracks was not sufficient to determine the residence time of any slower-dissociating population with confidence. Thus, consistent with the fact that only some of the auto-phosphorylation sites are mutated in this allele, Pom1^3A^ molecules bind the membrane longer than Pom1^WT^, but not as long as Pom1^KD^. The distribution of diffusion coefficients was also intermediate between Pom1^WT^ and Pom1^KD^ (Figure 2B). The thresholds defined above similarly showed a lower proportion of fast-diffusing molecules than Pom1^WT^ but a higher proportion than Pom1^KD^ (Figure 2C-D, Figure 2 – supplements 1A-C and 2). These data are consistent with auto-phosphorylation promoting Pom1 detachment from the plasma membrane.

To assess phosphorylation-dependent gradient shape changes in a simplified system containing a single membrane-binding site, we used the Pom1^MB1*^ mutant. This mutant binds the membrane solely with its amphipathic helix, which contains only phospho-sites 4 and 5. Consistent with poorer membrane binding, Pom1^MB1*^ gradient profiles showed decreased intensity compared to wildtype, at both cell poles and cell sides (Figure 4D). Mutagenesis of phospho-site 5, generating Pom1^MB1*-1A^, led to further gradient flattening with decreased Pom1 intensity at cell tips and increase at cell middle (Figure 4D). Thus, autophosphorylation regulates each of the two membrane-binding sites.

The progressive increase in medial cortical Pom1 levels in the phospho-site allelic series is consistent with the previously proposed idea that sequential phosphorylation events provide a timer function for Pom1 diffusion from cell poles. Additionally, non-phosphorylated Pom1 alleles may directly bind the membrane at the cell sides. To test the second scenario, we monitored the localization of the allelic series in *tea4Δ* cells, which lacks the phosphatase regulatory subunit (Alvarez-Tabares et al., 2007; Hachet et al., 2011). In *tea4Δ* cells, all Pom1 phospho-mutants bound the cortex nearly uniformly (Figure 4A, bottom row), with levels that increased with the number of phospho-site mutations (Figure 4B-C, right panels). Again, Pom1^5A^ cortical levels were indistinguishable from Pom1^6A^, but a little lower then Pom1^KD^, for unknown reasons. These data are in agreement with the idea that each phosphorylation event progressively lowers membrane affinity to reduce Pom1 binding in the medial region. We note that changes in Pom1 distribution in the phospho-site mutants are not due to a change in Pom1 protein concentration, as verified by Western blot analysis (Figure 4 – supplement 1B). Comparing Pom1 medial cortical levels in WT and *tea4Δ* cells showed significantly lower amounts for Pom1 and Pom1^1A^ in *tea4Δ*, but higher or similar levels for Pom1^2A^, Pom1^3A^ and Pom1^5A^. This suggests that the flatter gradients observed in the Pom1 phospho-mutants are not only due to a reduction of Pom1 detachment from the membrane, allowing lateral diffusion over a longer distance, but also to an increase in Pom1 attachment to the membrane at mid-cell. Thus, Pom1 auto-phosphorylation both favours its detachment from, and prevents its attachment to, the membrane.

### Pom1 phospho-mutants have reduced gradient shape robustness

A key feature of the Pom1 gradient is its robustness to variations within the system. Previous work showed that variability in Pom1 concentration at cell poles is counteracted by varying the gradient decay length, which leads to a strong negative correlation between decay length and Pom1 amplitude at the pole (Hersch et al., 2015; Saunders et al., 2012). Pom1 clustering and Pom1 inter-molecular phosphorylation have both been theoretically proposed as source for this correction, though we so far lack experimental evidence. We assessed the contribution of the multi-phosphorylation reaction in gradient buffering by plotting the correlation between decay length and amplitude at the pole in Pom1 phospho-mutants that retain significant pole enrichment. Pom1^1A^, Pom1^2A^ and Pom1^3A^ all showed a loss of negative correlation of decay length to Pom1 concentration at the cell tip (Figure 4E). Thus, these Pom1 mutants poorly correct intrinsic variations of Pom1 concentrations at the cell tips. Another evidence for buffering comes from the steep decrease in the coefficient of variation of Pom1^WT^ from the cell tip to the cell middle (Hersch et al., 2015). While we confirm this observation with our current data set, the decrease in the coefficient of variation at cell tips and cell sides is much smaller for Pom1^1A^, Pom1^2A^ and Pom1^3A^ (from 9.6 for Pom1^WT^ to 6.6 for Pom1^1A^, 5.8 for Pom1^2A^, and 4.2 for Pom1^3A^; Figure 4F, Figure 4 – supplement 1C). We conclude that the Pom1 phosphorylation cycle directly contributes to Pom1 gradient shape robustness.

### Pom1 at the mid-cell cortex controls cell length at division

One important physiological role of Pom1 is to set cell size at division by negatively regulating the SAD-family kinase Cdr2, which forms stable cortical clusters at mid-cell (Martin and Berthelot-Grosjean, 2009; Moseley et al., 2009). Consistent with previous reports that Pom1^6A^ and Pom1 overexpression lead to cell size increase at division (Hachet et al., 2011; Martin and Berthelot-Grosjean, 2009; Moseley et al., 2009), we observed a gradual increase in cell length at division in mutants of the phospho-site allelic series (Figure 5A). Cell length at division was strongly correlated with the medial cortical Pom1 levels (Figure 5B), consistent with the idea that medial Pom1 levels set the cell division size. To evaluate the relative contributions of cytosolic and cortical Pom1 in Cdr2 inhibition, independently of gradient formation, we obtained the same measurements in *tea4Δ* cells: the progressive increase in cortical Pom1 in Pom1 phospho-mutants also correlated with an increase of cell length in *tea4Δ* background (Figure 5A-B). We note that *tea4Δ* cell lengths were slightly longer than WT for *pom1* alleles containing up to 3 phosphosite mutations. By contrast, the correlations between cell length and Pom1 levels at cell poles ran in opposite directions in WT and *tea4Δ* cells (Figure 5 – supplement 1A). We conclude that the medial cortical pool of Pom1 is the relevant pool for cell size regulation.

**Figure 5.**
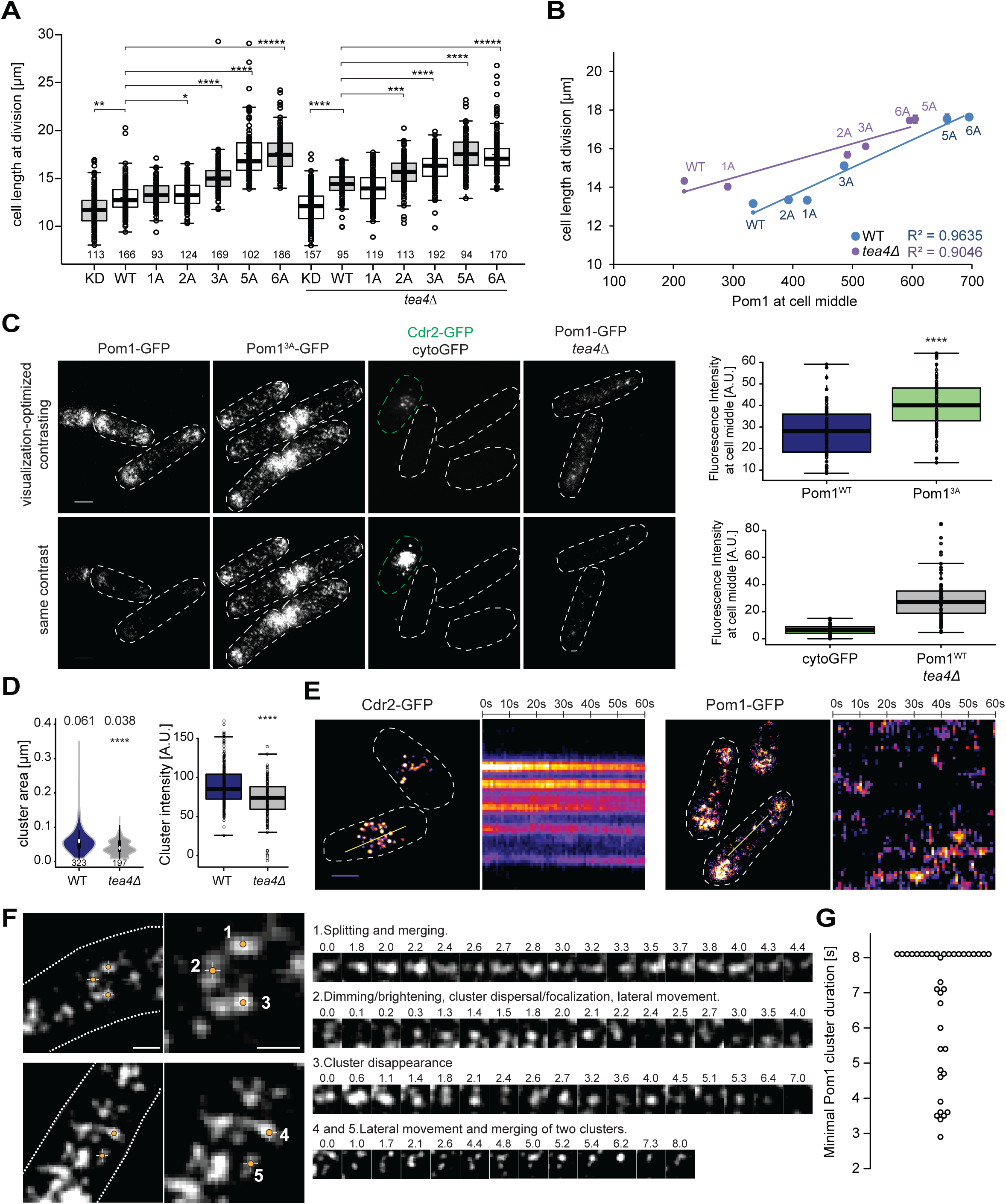
Cortical Pom1 at mid-cell regulates cell size. **A.** Mean cell length at division of the *pom1* phospho-mutant allelic series in otherwise wild type and *tea4Δ* background with number of quantified cells indicated, *, p=10^−2^, **, p=10^−7^, ***, p=10^−10^, ****, p≤10^−20^, *****, p=10^−40^. **B.** Correlation plot of cell length at division (values from panel A) versus Pom1 intensity at cell middle (values from Figure 4C). Error bars are standard error. **C.** TIRF imaging of Pom1, shown with identical acquisition and contrasting parameters (bottom) and adjusted contrast (top). Quantification of intensities is shown on the right (strains imaged the same day plotted together). Pom1^3A^-GFP shows increased cortical levels compared to Pom1^WT^-GFP (p=10^−9^). Note that cytoGFP expressed from the *pom1* promoter strain was mixed with a Cdr2-GFP strain to identify the TIRF focal plane (green dotted line). Scale bar 2.5μm. **D.** Mean cluster area (left) and average cluster intensity (right) for individual Pom1 clusters from wild type versus *tea4Δ* background, n=25 cells from 2 individual experiments. Number of clusters indicated below violin plot. The mean area is shown at the top, ****, p=10^−15^. **E.** Localization of Cdr2-GFP and Pom1-GFP by TIRF microscopy. Left panels show a snapshot of time point 0 from a time series imaged every second for 60s. Right panels show kymographs along the dotted line. Scale bar 2.5μm. **F.** Examples of Pom1-GFP cluster behaviours in TIRF, taken from two cells imaged every 100ms over 8s. Line 1: Cluster splitting (3.0s and 4.4s) and merging (3.2s). Line 2: Cluster fluorescence fluctuations (down at 0.1s, 2.0s, 2.4s; up at 0.2s, 2.1s, 2.5s). This cluster also exhibits clear lateral movement. Line 3: Cluster fluorescence fluctuation (down at 2.4s and 3.6s; up at 2.6s and 4.0s) and disappearance (7.0s). Line 4: Merging of two clusters (at 6.2s). The bottom cluster exhibits clear lateral movement. Scale bars: 1µm. **G.** Minimal lifetimes of Pom1 clusters in seconds. Note that the lifetime of all but 2 clusters is underestimated, as they either existed at the start or end of the timelapse, or both, n=38 clusters from 12 cells.

Because Pom1 clusters rather than individual molecules may shape the gradient (see Figure 1 above), and inspired by recent work showing visits of Cdr2 nodes by the downstream Wee1 kinase (Allard et al., 2018; Gerganova and Martin, 2018), we turned to live cell TIRF imaging. To test the method’s sensitivity and selectivity for cortical signals, we first compared the fluorescence levels of Pom1^WT^ and Pom1^3A^ phospho-mutant, which reproduced the increased Pom1^3A^ medial cortical localization seen by confocal microscopy (Figure 5C). We then imaged two cytosolic proteins. First, cytosolic GFP, expressed under the *pom1* promoter, could not be detected by TIRF though it was seen by epifluorescence, confirming that the evanescent field detects only cortical molecules (Figure 5C, Figure 5 – supplement 1B). By contrast, Pom1-GFP in *tea4Δ* cells, which by confocal microscopy is not detected at the cortex (see Figure 4A; (Hachet et al., 2011; Kokkoris et al., 2014; Padte et al., 2006)), revealed a cortical signal by TIRF imaging (Figure 5C). In *tea4Δ*, Pom1 formed cortical clusters, though cluster number, size and intensity were lower than in wild type (Figure 5D), which likely explains why the cortical Pom1^WT^ signal in *tea4Δ* is virtually indistinguishable from the cytosolic signal by confocal microscopy. Thus, very transient encounters of Pom1 at Cdr2 nodes at the cortex, rather than a fully cytosolic Pom1, likely account for the longer size of *tea4Δ* cells noted above. We conclude that TIRF provides a highly specific and sensitive imaging setup for Pom1 clusters at the yeast cortex.

In TIRF timelapse imaging, Cdr2-GFP formed stable nodes at mid-cell, which did not move substantially over 60s (Allard et al., 2018), whereas Pom1-GFP clusters were highly dynamic (Figure 5E). Fast 100ms acquisition intervals were used to monitor Pom1 clusters, which exhibited an array of dynamics with clusters moving laterally, splitting or merging. Figure 5F and Supplementary videos 1-2 provide representative examples. Example 1 shows an initial large cluster that splits in two at 3.0s, remerges, and splits again at 4.4s. Clusters 4 and 5 provide a second example of a small cluster moving laterally and merging with a larger one. Another common cluster dynamic is exemplified by clusters exhibiting dimming and/or dispersal of signal, followed by an immediate increase in fluorescence. This behaviour can be seen in example 2 between times 0.0s and 0.3s, 1.8s and 2.1s, and 2.2s and 2.5s, the latter one dimming to only a few detectable fluorescent pixels. Similarly, example 3 dims and regains fluorescence between 2.1s and 2.6s, and 3.2s and 4.0s. It then completely disappears at 7.0s. There were also many instances where the Pom1 signal was too fluid to unambiguously follow individual clusters over time. These fluctuations in the fluorescence signal of individual clusters indicate that clusters often recombine and can gain and loose individual Pom1 molecules over time.

To estimate the lifetime of individual clusters, we followed 38 Pom1 clusters in 12 cells. The cluster lifetime from appearance to disappearance or splitting of the cluster ranged to over 8s, the length of the imaging timeframe (Figure 5G). This value is very likely to be underestimated as the longest-lived clusters were present from start to end of the time-lapse imaging. These values are higher than previous measurements obtained by confocal microscopy (Saunders et al., 2012), probably because of the higher sensitivity of TIRF imaging.

They are also substantially longer than the binding time obtained by PALM imaging for individual molecules (around 1 to 3s, see Figure 1). These observations further support the idea that individual Pom1 molecules turn over within single clusters. Therefore, we propose that Pom1 clusters are the functional units shaping the gradient, as their longer residence time would permit diffusion from the cell pole all the way to the zone of action at mid-cell.

### Pom1 overlaps more with Cdr2 in short than long cells

To investigate the Pom1-Cdr2 interaction at the cortex, we acquired Pom1 TIRF images at 1s interval for 60s and took snapshot TIRF images of Cdr2 at the start and end of the imaging period in a dual tagged Pom1-GFP Cdr2-tdTomato strain (Figure 6A). The Cdr2 snapshots provided the location of clusters to which we mapped individual regions of interest (ROIs), in which we quantified Pom1 intensity in the GFP channel. Indeed, we were able to observe Pom1 encounters of the Cdr2 clusters of various duration within the 60 seconds imaging period (Figure 6A-B). To distinguish whether these encounters are targeted visits or due to random collisions of Pom1 clusters, we shifted the same size ROIs away from, but in the immediate vicinity of, Cdr2 nodes. We observed a very similar pattern for Pom1 in the non-node-associated ROIs, which suggests that laterally moving Pom1 clusters randomly collide into Cdr2 nodes (Figure 6C), distinct from the targeted visits from the cytosol reported for Wee1 (Allard et al., 2018).

**Figure 6.**
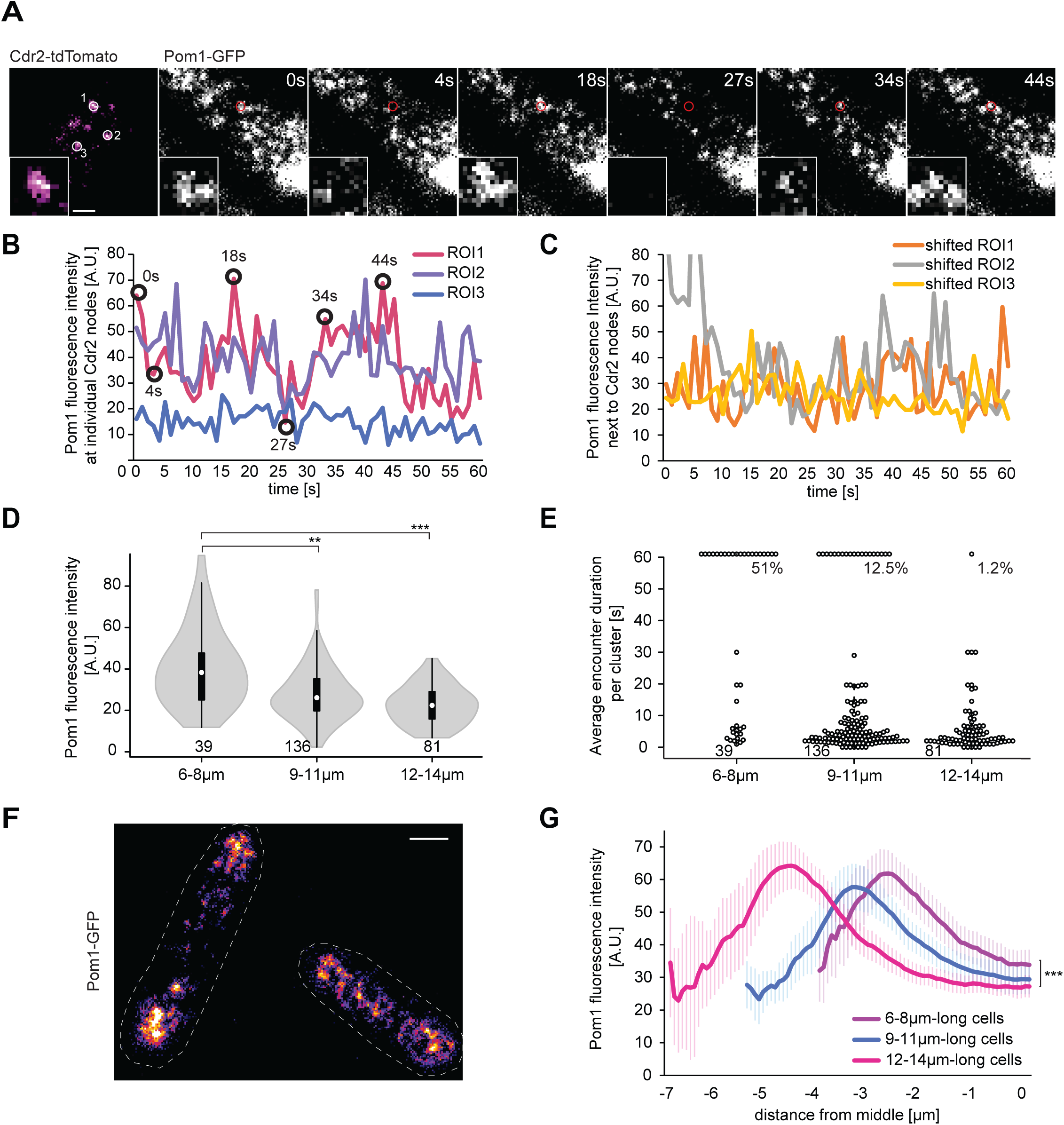
Pom1 levels at Cdr2 nodes are higher in short than long cells. **A.** TIRF images of Cdr2-tdTomato and Pom1-GFP in a dual tagged strain. ROI (1, 2 and 3) were selected around Cdr2 clusters and fluorescence intensity measured in the GFP channel (ROI 1 marked in red). Scale bar 1μm. **B.** Pom1-GFP fluorescence intensity in the three ROIs marked in panel A. Circled time points correspond to snapshots in panel A. **C.** Pom1-GFP fluorescence intensity in ROIs shifted to the immediate vicinity of the ROIs marked in A. **D.** Average fluorescence intensity of Pom1-GFP at Cdr2 nodes measured as in (A-B) over a 60s imaging period, sorted by cell length.***, p≤10^−7^. **E.** Duration of individual Pom1 encounters with a Cdr2 node for cells of sorted length. This is defined as the length of time the Pom1-GFP signal is over 20 arbitrary fluorescence units. In longer cells, the proportion of clusters with continuous (60s) Pom1 presence decreases, while the number of shorter encounters increases. For (D-E), that data was collected from a single experiment: 39 clusters from 6 6-8μm long cells, 136 clusters from 21 9-11μm long cells, and 81 clusters from 13 12-14μm long cells. Experiment duplicates presented in Figure 6 – supplement 1. **F.** Localization of Pom1-GFP in TIRF in cells of different lengths (10μm and 13.5μm). Scale bar 2.5µm. **G.** Global cortical Pom1 levels measured from TIRF imaging. The gradients are aligned to the cell geometric middle and sorted by cell length. Error bars: standard error between experiments (N=3); in total 62 gradients from 6-8μm cells, 248 gradients from 9-11μm cells and 128 gradients from 12-14μm cells. Statistics on Pom1 medial levels performed for the last 1.5μm of the gradient tail between short (6-8μm) and intermediate cells (9-11μm), p=10^−3^, and between short and long cells (12-14μm), p=10^−6^.

Remarkably, when we clustered the data according to cell length, the average Pom1 intensity at all measured Cdr2 nodes was significantly higher in short (6 to 8µm) than long (12 to 14µm) cells in three independent experimental repeats (Figure 6D, Figure 6 – supplement 1A-B). The values for cells of intermediate length (9 to 11µm) were more variable, probably depending on the average length of these cells. We observed a similar pattern in the duration of Pom1-Cdr2 encounters, which we defined as the length of time the Pom1-GFP value at a Cdr2 node remained above a defined fluorescence threshold: In short cells, a higher proportion of Cdr2 clusters were continuously occupied by Pom1 over the 60s imaging period (Figure 6E, Figure 6 – supplement 1C). Consistent with the observation that Pom1 behaviour is similar at and in the vicinity of Cdr2 nodes, measuring the total medial Pom1 TIRF signal gave similar results: Pom1 levels were higher in the middle of short than long cells (Figure 6F-G). Thus, the medial cortical Pom1 levels, which are those critical for Cdr2 regulation, decrease as cells grow, consistent with progressive cell length-dependent relief of Pom1 inhibition.

## Discussion

Investigations on Pom1 concentration gradients have provided two main lines of discussion, one concerning the mode of gradient buffering and another concerning the function of the gradient as a measure of cell length to control the timing of mitotic commitment. In this work we provide evidence for unified models on these two subjects. We confirm experimentally that the buffering of the Pom1 concentration gradient relies on its multi-phosphorylation reaction and establish that Pom1 molecules turn over within membrane-bound clusters. The claim of Pom1 gradients serving as a measure of cell length was originally tempered by further investigations that did not detect differences in Pom1 levels at mid-cell. By specifically imaging the relevant pool of Pom1 at the mid-cell cortex, we now unequivocally show that Pom1 gradients formed by diffusion of Pom1 clusters reach the middle of short but not long cells.

### Pom1 clusters as diffusion unit to shape gradients

The Pom1 gradient forms upon local dephosphorylation of Pom1 at the cell pole, promoting Pom1 membrane binding (Hachet et al., 2011). Previous modelling work demonstrated that the multi-phosphorylation reaction Pom1 undergoes prior to its membrane detachment could provide a timer function for Pom1 lateral diffusion and serve as a buffering mechanism for the gradient (Hersch et al., 2015). An alternative model proposed that differential diffusion coefficients of Pom1 clusters of varying sizes underlie the buffering of the gradient with the largest clusters causing a “traffic jam” event at the cell tip, which is relieved by the progressive fractionation and the subsequent increase in diffusion coefficients of smaller clusters away from the gradient source (Saunders et al., 2012). The data presented here integrates these two mechanisms in a single model for Pom1 gradient formation. Single molecule measurements by PALM imaging show that the gradient is comprised of molecules with a wide distribution of diffusion coefficients. Their dissociation dynamics also revealed at least two populations with distinct binding times. Note that the fast-dissociating population is not identical to the fast-diffusing one. However, there is an overall inverse correlation between binding times and diffusion rates, such that fast-diffusing molecules are also binding the membrane only for a short time. These may represent individual Pom1 molecules not associated with a cluster or in small clusters, and are a minority of all Pom1 molecules. The slower-diffusing molecules may be part of larger clusters. Many of these molecules detach from the membrane slower, but a large pool also exhibits fast dissociation, perhaps indicating peripheral association with the cluster. Importantly, the diffusion coefficients and binding times reveal that individual molecules (in a cluster or not) will only travel a maximum distance of 0.8 µm before detaching from the plasma membrane. Thus, a Pom1 molecule binding at the cell tip upon dephosphorylation by the Tea4-PP1 complex travels only a short distance and does not reach the cell middle.

By contrast, Pom1 clusters as a whole are longer-lived at the plasma membrane. Clusters appeared very dynamic, changing intensity over time, splitting, dispersing and fusing again. Importantly, TIRF measurements easily identified clusters lifetimes of over 8 s. This longer lifetime of clusters at the plasma membrane indicates that individual molecules exchange within a cluster. Specifically, this suggests that existing clusters must be able to bind Pom1 molecules directly from the cytosol. In TIRF imaging, we indeed found examples of isolated clusters whose fluorescence intensity diminished before increasing again, providing support for this idea. These longer lifetimes are consistent with clusters being able to form at the cell pole and diffuse laterally all the way to mid-cell, at least in short 7-8 µm cells. Thus, we propose that Pom1 clusters are the functional units that shape the Pom1 graded distribution.

### Pom1 phosphorylation shapes the gradients

The phospho-site mutant allelic series shows that Pom1 distribution critically depends on auto-phosphorylation. Indeed, progressive alanine substitution at up to 5 auto-phosphorylation sites causes progressive flattening of the Pom1 graded distribution, with apparently additive contribution of each phosphorylation event. Because the 5 phosphorylation sites all map very close to each other within two adjacent membrane-binding motifs, it is likely that they have to be phosphorylated sequentially, extending the time frame between the dephosphorylation event taking place at the cell pole and the full auto-phosphorylation, promoting membrane detachment.

Each phosphorylation event reduces the affinity to the plasma membrane, likely affecting both k_off_ and k_on_. Indeed, our PALM data shows that the k_off_ of single Pom1 molecules is modulated by their phosphorylation status: Pom1^KD^, which is not phosphorylated binds the plasma membrane longer than the partly dephosphorylated Pom1^3A^, which itself binds longer than Pom1^WT^. Similarly, alanine-substitutions of phospho-sites led to a progressive increase in the membrane-associated Pom1 fraction in *tea4Δ* cells, with a fold-change larger than the one measured for the k_off_. This suggests that, although not directly measured, the k_on_ of Pom1 to the plasma membrane is also modulated by phosphorylation, which decreases the membrane association rate.

Phosphorylation may also modulate cluster formation. Clusters formed in all studied Pom1 mutant alleles and exhibited a similar wide range of diffusion rates in Pom1^WT^, Pom1^3A^ and Pom1^KD^, suggesting – if we make the validated assumption that cluster size influences diffusion rate (Saunders et al., 2012) – a similar range of cluster sizes. Thus, dephosphorylated Pom1 efficiently forms clusters. Phosphorylated Pom1 may form clusters less efficiently, as Pom1^WT^ forms smaller clusters in *tea4Δ* cells. We note that it is unclear whether Pom1 is fully phosphorylated in this mutant, or whether it may be inefficiently dephosphorylated by the still present PP1 catalytic subunit lacking the Tea4 regulatory subunit. Previous data had indeed shown that Pom1 dephosphorylation does not strictly require Tea4 (Kokkoris et al., 2014). In either case, the data indicate that dephosphorylation at the cell pole also promotes clustering, which may be further favoured by the direct binding with Tea4 (Hachet et al., 2011). We note that it may be impossible to fully dissociate cluster formation from membrane binding.

The comparison of Pom1^WT^ and Pom1^KD^ reveals three important differences. First, the diffusion rates of Pom1^KD^ at cell sides were similar to those of Pom1^WT^ at cell poles, consistent with local dephosphorylation of Pom1 at this location. Second, on cell sides the fast pool of Pom1^WT^ molecules was more abundant and diffused faster than Pom1^KD^. This suggests that, by reducing the electrostatic interaction with neighbouring phospholipids, auto-phosphorylation promotes faster mobility of individual Pom1 molecules. Finally, the proportion of single Pom1 molecules increased with distance from cell poles in Pom1^WT^ but not Pom1^KD^. These observations are consistent with progressive phosphorylation-induced dissociation of Pom1 from the membrane and/or clusters.

From these data, we propose a revised model on how the specific gradient shape is achieved. Localized Tea4-PP1 phosphatase activity at cell poles dephosphorylates Pom1, revealing two membrane-binding domains – an amphipathic helix and an adjacent positively charged region – both of which permit the association of Pom1 with the plasma membrane. Pom1 dephosphorylation also favours the formation of clusters at the membrane, which helps carry Pom1 over a longer distance. Within a cluster, individual Pom1 molecules have a short lifetime of 1 to 3 seconds on average, but new Pom1 molecules join from the cytosol. As clusters split and merge, this permits the transport of single molecules from cluster to cluster. The clusters may diffuse at different rates, either because they are of different sizes and therefore contain different numbers of membrane-binding sites, or because each membrane binding site may be differently phosphorylated and thus bind the membrane with distinct affinity. Indeed, as Pom1 is active on itself, aging clusters become more phosphorylated, which promotes dissociation from the cluster and from the membrane to single molecules. Even though single molecules are present at the membrane a long distance from the poles, their binding is short-lived, detaching from the membrane within 1 second. Thus, clusters act as a diffusion unit, whose regenerating capacity is reduced via progressive auto-phosphorylation with time/distance from the cell pole

### Pom1 medial cortical levels control mitotic commitment and vary with cell size

One physiological function of Pom1 is to prevent the activation of Cdr2 kinase. Our data clearly establish that the meaningful levels of Pom1 are those at the mid-cell cortex, where Cdr2 forms nodes. Indeed, we see a strong correlation between cell size at division (the phenotypic outcome of Cdr2 regulation) and Pom1 medial levels in both WT and *tea4Δ* cells. By contrast, the correlations between cell pole levels of Pom1 and cell length is inverse in WT and *tea4Δ* cells, excluding the alternative model that Pom1 may act on Cdr2 at cell poles (Bhatia et al., 2013). These data agree with the extended size of cells overexpressing Pom1 or mis-targeting it to the medial cortex (Martin and Berthelot-Grosjean, 2009; Moseley et al., 2009). They also agree with the strong cell lengthening effect of naturally re-distributed Pom1 upon glucose starvation (Kelkar and Martin, 2015). Our new observation that *tea4Δ* cells have significant cortical Pom1 signal also fits with the idea that Pom1 remains active on Cdr2 in this mutant as manifested by the longer size of *tea4Δ* than *pom1Δ* (this work and (Martin and Berthelot-Grosjean, 2009)). In this mutant, abundant cytosolic Pom1 likely allows substantial stochastic encounter with, and binding to, the plasma membrane. We note that the phosphomutant Pom1 alleles did not cause noticeable changes in the position of the division site. This is in agreement with the finding that the two functions of Pom1 in controlling positioning and timing of division are separable (Bhatia et al., 2013). This also indicates that a graded Pom1 distribution is less important for division site placement, consistent with observations in *S*. *japonicus* (Kinnaer et al., 2019).

Although Pom1 mid-cell levels clearly correlate with cell size at division in various mutant and environmental conditions, a key unresolved and debated question has been whether Pom1 levels vary during the growth of a single cell, and therefore whether Pom1 may contribute to cell size homeostasis. The higher sensitivity and specificity of TIRF imaging now allows to answer this question unequivocally: there is more Pom1 at the mid-cell cortex of small than long cells. We considered the possibility that this higher concentration may be a simple consequence of cell extension. However, a fission yeast cell doubling in length increases its volume about 1.1-fold more than its surface. Thus, considering a constant Pom1 concentration and an invariant average gradient shape (Bhatia et al., 2013; Saunders et al., 2012), a higher number of Pom1 molecules have less membrane space at their disposal, which would lead to higher Pom1 concentration at mid-cell in longer cells. Because we observe the opposite, the increased mid-cell levels of Pom1 in short cells must be due to Pom1 gradients extending into mid-cell of small but not long cells. The diffusion rates and cluster lifetimes we have measured are consistent with this scenario. These may be tuned to allow Pom1 diffusion over 3-4µm distance, sufficient to reach the middle of short, but not long cell, which underlies the difference in Pom1 mid-cell cortical levels in cells of different sizes. Thus, the graded Pom1 distribution is able to convey information in a manner dependent on distance from the source.

These findings poise Pom1 to function as a sensor of cell dimension that provides more inhibition on Cdr2 in short than long cells. This proposition is consistent with biochemical data that Cdr2 activating phosphorylation, which Pom1 counteracts, increases with cell growth (Deng et al., 2014) and that Cdr2 is more active in long than short cells, as measured by the number of visits by the Wee1 kinase (Allard et al., 2018). It is also consistent with Pom1 function being exquisitely dose-dependent, both in terms of global levels (Martin and Berthelot-Grosjean, 2009; Moseley et al., 2009) and specifically at the mid-cell cortex (this work). Pom1 may not be the only sensor protein in the pathway: Cdr2 was also proposed to monitor cell surface area through a dynamic exchange of molecules between the cytoplasm and the plasma membrane to form medial nodes whose numbers increase with cell growth (Facchetti G et al., 2019; Pan et al., 2014). Thus, cell growth, by both promoting a decrease in Pom1 levels and an increase in the number of Cdr2 nodes at mid-cell, enhances the activation of Cdr2 in long cells. As mutants in both Pom1 and Cdr2 retain cell size homeostasis, cells likely have secondary sizer mechanisms, perhaps monitoring different geometrical quantities (Facchetti G et al., 2019; Wood and Nurse, 2013).

## Materials and methods

### Yeast strains, Media, and Genetic Approaches

Standard methods of *S*. *pombe* culturing and genetic manipulations were used. For PALM imaging, *S*. *pombe* cells were grown in rich yeast extract (YE) medium and imaged during the exponential growth at an OD_600_ comprised between 0.4 and 0.7. All live-cell imaging was performed on YE 2%-agarose pads. For all other imaging experiments strains were grown at 25°C in fully supplemented synthetic Edinburgh minimal medium (EMM). A complete list of all used strains is provided in Table S1.

For the truncation analysis (Fig. 3D), Pom1 fragments were amplified by PCR and cloned under control of the *pom1* promoter in single integration vectors (kind gifts from Dr. Aleksandar Vjestica). The vectors were linearized and integrated at the *ura4* locus in wild type and *pom1Δ* strains. A list of plasmids used in this study is provided in Table S2. Plasmids maps are available upon request. The I494N mutation was introduced by site-directed mutagenesis. To generate the MB1* allele, we made use of a native restriction enzyme site (BglII) at aa426 to replace fragment 426-510 with one in which aa429-436 within the conserved region (MB1) were replaced by 7 alanines. This mutated fragment was amplified with a forward primer annealing from aa436 onward, and carrying an overhang for the 7 alanines and the BglII restriction site. Primer sequences used for mutagenesis are listed in Table S3. To generate *pom1* alleles at the native genomic locus (Fig 3E), pSM2142 (containing *pom1* promoter, ORF fused in frame to GFP, kanMX and *pom1* 3’UTR) was used as a backbone for site-directed mutagenesis, except for the *pom1*^*MB1\**^ allele, for which the 7 residues were changed to alanine in 3 rounds of site-directed mutagenesis. To generate the MB1* I494N 5PxxP* mutant, a sequence including the sequence coding for aa81-524 (BspEI to MluI) containing all mutations (as above and in (Hachet et al., 2011)) except PxxP sites 4 and 5 was ordered as a synthetic gBlocks® from LubioScience GmbH and introduced in replacement of the wildtype fragment in plasmid pSM2142. PxxP sites 4 and 5 were then introduced by site-directed mutagenesis. The plasmids were digested and transformed into a *pom1Δ∷ura4+* background.

The phosphosite mutant allelic series (Fig. 4A) was generated through site-directed mutagenesis of the sites on a pREP41 plasmid, carrying the full length Pom1 sequence (pSM738). The mutagenized plasmid was digested and transformed in a strain in which *ura4+* replaced the *pom1* sequence coding for aa400-1006 (*pom1Δ(aa400-aa1006)∷ura4+*). Integrants were selected on 5-FOA. Each *pom1* allele was subsequently tagged with GFP-kanMX through transformation with linearized pSM1731, a plasmid containing the end of *pom1* ORF without stop codon fused in frame to GFP, the kanMX selection marker and *pom1* 3’UTR. For the Pom1^MB1*-1A(5)^ mutant, phosphosite 5 was mutagenized through site-directed mutagenesis on pSM2237 (Pom1^MB1*^-GFP) to generate pSM2264, which was digested and transformed into a *pom1Δ∷ura4+* strain as above.

All generated strains were verified via sequencing.

Pom1-mEos3.2, Pom1^KD^-mEos3.2, and Pom1^3A^-mEos3.2 strains were generated for PALM microscopy using a standard PCR-based approach (Bähler et al., 1998; Laplante et al., 2016) and verified by PCR.

BH-search prediction performed at https://hpcwebapps.cit.nih.gov/bhsearch/ with window size for residue averaging of 15 amino acids and values for amino acids set to standard BH parameters.

### Microsized-hole preparation for vertical immobilization of *S*. *pombe* cells

For cell pole imaging, *S*. *pombe* cells were vertically immobilized on a YE-2% agarose pad obtained by imprinting on an epoxy resin mold containing an array of micro-pillars (Wang and Tran, 2014) with diameter of 6 μm and height of 20 μm (Figure 1 – supplement 1).

SU-8 photolithography and PDMS lithography were performed at the EPFL Center of Microtechnology. SU-8 (GM1060 Gersteltec Sarl) was spin-coated onto a silicon wafer with a thickness of 20 μm and then baked at 95°C for 40min. The wafer was then gradually cooled from 95°C to 30°C during 30 min. SU-8 polymerization was induced by exposure to 350nm light for 10 s through a quartz mask containing disk patterns of 6 μm diameter. Post-photolithography baking was performed at 95°C for 40 min, followed by gradual cooling for 30 min. SU-8 was then developed with manufacturer-provided SU-8 developer and cleaned with isopropanol and dried with compressed air. The SU-8 substrate was hardened by baking at 150°C for 30 min.

PDMS (Sylgard 184 silicone base, Sylgard curing agent) was then poured onto the surface of the SU-8 mold, which was pre-treated with trimethylchlorosilane vapor (TMCS 33014 from Sigma) to render it non-stick. The PDMS was then polymerized during at least 2 h at 80°C. The resulting PDMS mold was then used to create the final epoxy resin mold. Epoxy resin R123 bisphenol and epoxy hardener R614 (Soloplast Vosschemie) were poured onto the PDMS substrate and polymerized for 24 h.

### Super-resolution microscopy for Pom1 dissociation and diffusion dynamics

*S*. *pombe* cells were genetically modified to express the photoactivable fluorophore mEos3.2 fused to Pom1 protein at physiological level (Laplante et al., 2016).

To measure diffusion dynamics, imaging was performed on a previously described custom-built microscope (Holden et al., 2014). Cells were imaged in two channels: the fluorescence channel for precise tracking of single Pom1 molecules and the phase contrast channel for cell segmentation and determination of the membrane plane. In order to selectively image Pom1 at the membrane, we used astigmatic imaging to encode the axial position of single molecules (Huang et al., 2008). In a post processing step, we analysed single molecules which were at axial positions ranging from −200nm to +200nm (with 0 corresponding to the plasma membrane position), eliminating signal detected off the membrane. A z-calibration was performed by taking sequential images of 0.1 µm fluorescent beads (Invitrogen TetraSpeck), displacing the objective by 20 nm steps over an axial range of 1 µm. This calibration allows us to define the PSF widths in the x and y (lateral) directions as a function of the z (axial) displacement. The imaging was performed with an NA 1.49 oil immersion objective lens (Nikon), and fluorescence was detected using an Evolve 128 EMCCD camera (Photometrics) with a 20 ms integration time. Fluorescence was excited with a 560 nm laser (MPB VFL-P-300-560) using an irradiance of 4 kW/ cm^2^. Molecules of mEos3.2 were photoconverted using a 405 nm laser (Coherent OBIS) with an irradiance of ∼ 0-16 W/ cm^2^. To measure dissociation, Pom1-mEos3.2 molecules were photoconverted by a pulse of 405 nm and then imaged continuously (no time-lapse + 20 ms exposure time) or with the time-lapse sequences (time-lapse durations of 100+20ms or 200+20ms, Fig. 1c) with 560 nm light until complete bleaching of the photoconverted molecules before the cycle was repeated. Imaging of cells were performed until no more activation of Pom1-mEos3.2 was observed.

Brightfield illumination for phase contrast imaging was performed with a white LED (Thorlabs MCWHL2), passed through a green filter (Chroma ET525/50mc), focused into the back focal plane (BFP) of a condenser lens (Nikon MEL56100). This channel was equipped with a 1024×768 CMOS camera (The Imaging Source DMK 31BU03). Image acquisition was controlled through Micro-manager.

### PALM localization and image processing

The images were processed with a custom-made ImageJ plugin to segment the single molecule data. Briefly, this consisted of generating a maximum intensity projection (MIP) of all spots, dilating and binarizing the MIP to make a cellular mask and then multiplying this mask by the raw image data. This ensures signal exclusively within cells is analysed and eliminates spurious noise occasionally detected outside of a cell. 3D molecule localization was then performed with RapidSTORM 3.3 (Wolter et al., 2012) software, with a z calibration (PSF X and Y width versus Z) used as an input. Single molecules which had an SNR above 50 were localized. Subsequently, a wobble distortion was corrected using an open source Matlab script (Carlini et al., 2015); this eliminated lateral shifts in localizations above and below the focal plane. Before tracking, single molecules were filtered based on their integrated intensity and axial position to ensure that only molecules on the membrane were tracked. Localized molecules were then tracked using a Matlab-based routine based on (Crocker and Grier, 1996a): molecules belonged to the same track if they were within a 320 nm radius within consecutive frames. No gaps within tracks were permitted. The distance of each track’s first point was calculated relative to the cell pole, which was determined from the phase contrast image. First, the phase contrast channel was aligned to the fluorescence channel from images of 500 nm fluorescent beads (Invitrogen TetraSpeck) using a custom-written MATLAB program. Briefly, bead images were localized in 2D and then used to define a rigid transform using a custom MATLAB script. This transformation was then used to map the fluorescence channel onto the phase contrast channel. Second, the center-line of the cell were defined manually using MATLAB’s ‘imline’ function, with line extremities corresponding to the poles x,y positions.

### Dissociation and bleaching rate extraction and analysis

The tracking, dissociation and diffusion analysis was performed with custom Matlab script to extract and study the diffusion and dissociation dynamics.

The recorded molecule trajectories provide an apparent dissociation rate (*k*_*eff*_) which is the result of both the molecule photobleaching (*k*_*bleach*_) and the actual dissociation rate (*k*_*off*_) contributions. By performing time-lapse imaging at time-lapse periods (τ_*TL*_) of 20 ms, 120ms and 220ms we extracted the dissociation constant of Pom1 wide type or mutants.

First, the apparent dissociation rate (*k*_*eff*_) is extracted by fitting of the exponential distribution of the track lengths (*t*_*eff*_) for every time-lapse experiment (Figure 1D; Figure 1 – supplement 2A)

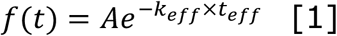

where the track length is defined as the number of sequential localization events (*n*) spatially separated by less than 321nm multiplied by the time-lapse period τ_*TL*_, *t*_*eff*_ = (*n* − 1) × τ_*TL*_, and τ_*TL*_ is the sum of the camera integration time (τ_*int*_– equal to 20ms in our experiment) and the time delays introduced within a pair of two consecutive images.

This distribution is the result of the sum of two independent Poisson processes: the photobleaching that occurs only under laser exposure (i.e. during τ_*int*_) and the dissociation of Pom1 that can occur any time during the τ_*TL*_ period.

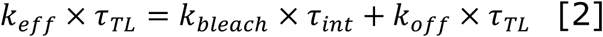

Thus, equation 2 can be rewritten as *y*(τ_*TL*_) = q + *m* × τ_*TL*_ where *k*_*off*_= *m* ands *k*_*bleach*_= q/τ_*int*_ are easily extracted from the slope and the intercept of the linear fit of *k*_*eff*_ × τ_*TL*_ versus τ_*TL*_ data. We performed a weighted linear fit where we assigned to each *k*_*eff*_ value a weight proportional to the S.D. extracted from the fit of the exponential distribution.

More details about the analysis and parameter results can be found in the Supplementary Notes and Table S4.

### Diffusion coefficient and dynamics analysis

The individual protein diffusion coefficient (*D*) is extracted from tracks containing more than five consecutive localizations without any gap between localizations using MSD analyser, a Matlab-based package (Tarantino et al., 2014). *D* is derived from the MSD distribution of a Brownian particle’s trajectories parameterized through the Einstein–Smoluchowsky equation *MSD* = 2*dD*Δ*t* where *d* is the number of dimensions of the trajectory data (*d* = 2 in this work, since we consider diffusion at the membrane) and Δ*t* is the time lag over which the MSD is measured. *D* can thus be extracted from the slope of linear fit of the first 25% of the mean MSD curve (Crocker and Grier, 1996b).

The equation 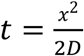 was used to derive diffusion distances from diffusion coefficients.

### Microscopy

Confocal microscopy (Fig. 3 and 4) was performed on an inverted DMI4000B Leica microscope equipped with an HCX Plan Apochromat 100x/1.46 NA oil objective and an UltraVIEW system (Perkin Elmer; including a real-time confocal scanning head CSU22 from Yokagawa Electric Corporation), solid state laser lines, and an electron-multiplying charge-coupled device camera (C9100, Hamamatsu Photonics). Medial section images were obtained at 400ms exposure time at 100% laser power for 5 consecutive time points at maximum speed with sum image projections used for quantification and figure preparation. For cell length measurements, cells were stained with calcofluor and imaged with a Leica epifluorescence microscope (60X magnification). TIRF microscopy (Fig. 5 and 6) was performed on a DeltaVision OMX SR imaging system, equipped with a 60x 1.49 NA TIRF oil objective (oil 1.514), an illumination pathway for ring-TIRF and a front illuminated sCMOS camera size 2560×2160 pxl (manufacturer PCO). Imaging settings were: 512×512 pxl field of view, 21ms exposure time, laser power of 20% with TIRF angles: 488nm at 86.9° and 568nm 83.2°. Samples were placed on a 0.17 +/-0.01 mm thick glass slide and imaged within 15 minutes. Imaging was performed in two modes: every second over a 60s imaging period and every 100ms over an 8 second imaging period. A widefield image of the medial plane of cells, expressing cytosolic GFP (kind gift from Dr. Magdalena Marek) mixed with a Cdr2-GFP strain (Fig. S5B) was taken as a single snapshot on the same imaging system, using light path setting conventional, 21ms exposure time, 20% laser power, after which the cortical plane of the same field of view was imaged in TIRF.

### Fluorescence quantifications

Cortical gradient profiles were quantified from confocal microscopy image sum projections by manually drawing a line along the cortex of the cell from the cell tip to the cell middle, generating 4 gradient profiles per cell in ImageJ (NIH). For the phosphosite mutant allelic series (Fig. 4B), 240 gradient profiles per strain were generated (n=3 experiments, 60 cells). 160 gradient profiles were generated for Pom1^MB1*^ and Pom1^MB1*-1A(5)^ (Fig. 4D) from n=2 experiments, 40 cells. aligned to gradient maximum intensity value at the cell tip, averaged per strain and plotted against distance from the cell pole. The three individual experiments are shown in Figure 4-S1. For the quantification of Pom1 intensity levels at cell middle, the average value over the medial-most 1.5μm of the gradient tails of each individual gradient was calculated and presented as boxplots (Fig. 4C). We developed a simple MatLab script to extract decay length (Suppl. Info 1). Briefly, all 240 gradient profiles were aligned to the maximum value at the cell pole and smoothed with a Gaussian filter as previously described (Hersch et al., 2015). The profiles were binned by 5% and the first 0.5μm of the profile values were deleted to avoid the effect of the gradient plateau at the cell pole. The decay length was obtained as the slope of the linear regression on the log of the binned average profiles and plotted against the log of the Pom1 amplitude at the pole. For the correlation plot between cell pole intensities to cell length (Fig S5A), the intensity at cell pole was calculated as the average values from the first 0.83μm of individual gradient profiles aligned to the maximum value at cell poles.

Cell length measurements (Fig. 5A) were performed on cells from 3 individual experiments (total number of quantified cells per strain is labelled on the figure), manually drawing a straight line at the detected calcofluor signal from cell tip to cell tip in ImageJ (NIH). Cells imaged at a medial plane. Individual lengths were recorded and presented as boxplots. For correlation plots to Pom1 levels at cell middle or at the cell pole, averages were calculated for each strain.

To quantify Pom1 cluster area and intensity (Fig. 5D), we drew ROIs around 323 individual clusters from 26 wild type cells and 197 individual clusters from 25 *tea4Δ* cells in ImageJ (NIH). The quantification was done on the image of the first time point from a 60s imaging period. For the quantification of cluster duration (Fig 5G), raw TIRF images were corrected for bleaching with the EMBL tool CorrectBleach and further smoothed in ImageJ. Clusters were then tracked individually for the duration of the 8 second imaging period. Fluorescence quantifications from TIRF imaging for global Pom1 levels (Fig. 6G) were performed by drawing a 45-pixel wide ROI across the entire detectable TIRF signal per cell. To generate two gradient profiles, one from each cell tip, the geometric middle of the cell was used for alignment. To measure intensities at cell middle, the values from the last 1.5μm of each gradient were averaged. The same method was used for the quantification shown in Fig 5C. For quantification of local Pom1 levels at Cdr2 nodes (Fig. 6B), a 9-pixel ROI was drawn around Cdr2-tdTomato nodes and used to detect signal in the Pom1-GFP channel. Control ROIs of the same size were shifted to the immediate vicinity of a Cdr2 node (Fig. 6C). Data corrections were done for background camera noise for each individual image and for bleaching. To estimate the average encounter duration, we recorded the length of time the Pom1 signal at a given node was above a defined fluorescence threshold. The threshold choice was instructed by the minimum detectable signal in that experiment. In all cases that threshold was higher than the fluorescence levels measured in the GFP channel using a strain that did not express GFP, in which Cdr2-tdTomato was used to identify the TIRF focal plane and the ROIs of interest. The subtracted exact arbitrary value is indicated in figure legends. Note that the difference observed between short and long cells was robust to changes in threshold choice.

### Western Blot

Yeast cultures were grown in YE medium at 30°C to OD_600_ = 0.8, collected by centrifugation at 3000rpm at 4°C for 5min and washed with 1x CXS buffer (50mM HEPES, pH 7.0, 0,20mM KCl, 1mM MgCl2, 2mM EDTA pH7.5 containing an anti-proteolitic tablet (Roche, Ref 05892791001). Lysates were obtained via mechanical breakage with acid-treated glass beads (Sigma), using a BeadBeater homogenizer for 10 repetitions at 4.5V of 30s on, 30s off on ice cycles. The samples were centrifuged at 10,000g for 20min at 4°C and extracts were recovered by pipetting into a new tube. Protein concentration was determined via spectroscopy using Bradford reagent. 270μg of proteins were loaded per sample on a 10% SDS-PAGE gel and transferred via wet Western blot transfer. 1° antibodies used: α-GFP 1:1000 dilution (Mouse, Roche, Cat.No. 11814460001), α-Tubulin (TAT1, Mouse, 1:5000 dilution), 2° antibodies α-mouse (HRP detection, Promega, W4021). The mean intensity quantification of 4 independent experiments is shown in Figure 4 – supplement 1B. Before averaging, values were corrected for the corresponding background and presented as a ratio of α-GFP to α-tubulin signal.

## Acknowledgements

We thank Dr Marileen Dogterom (TU Delft) and her group for providing the mask for photolithography for the fabrication of the micro-hole, Aster Vanhecke for assistance with processing of some of the tracking data, and Dr Aleksandar Vjestica and Dr Magdalena Marek for gifts of plasmids and strains. We thank Serge Pelet and members of the Martin lab for critical advice and comments on the manuscript. This work was supported by a Swiss National Science Foundation Sinergia Grant (CRSII3-160728) to SGM and SM.

## Figure legends

**Figure 1 – supplement 1.**
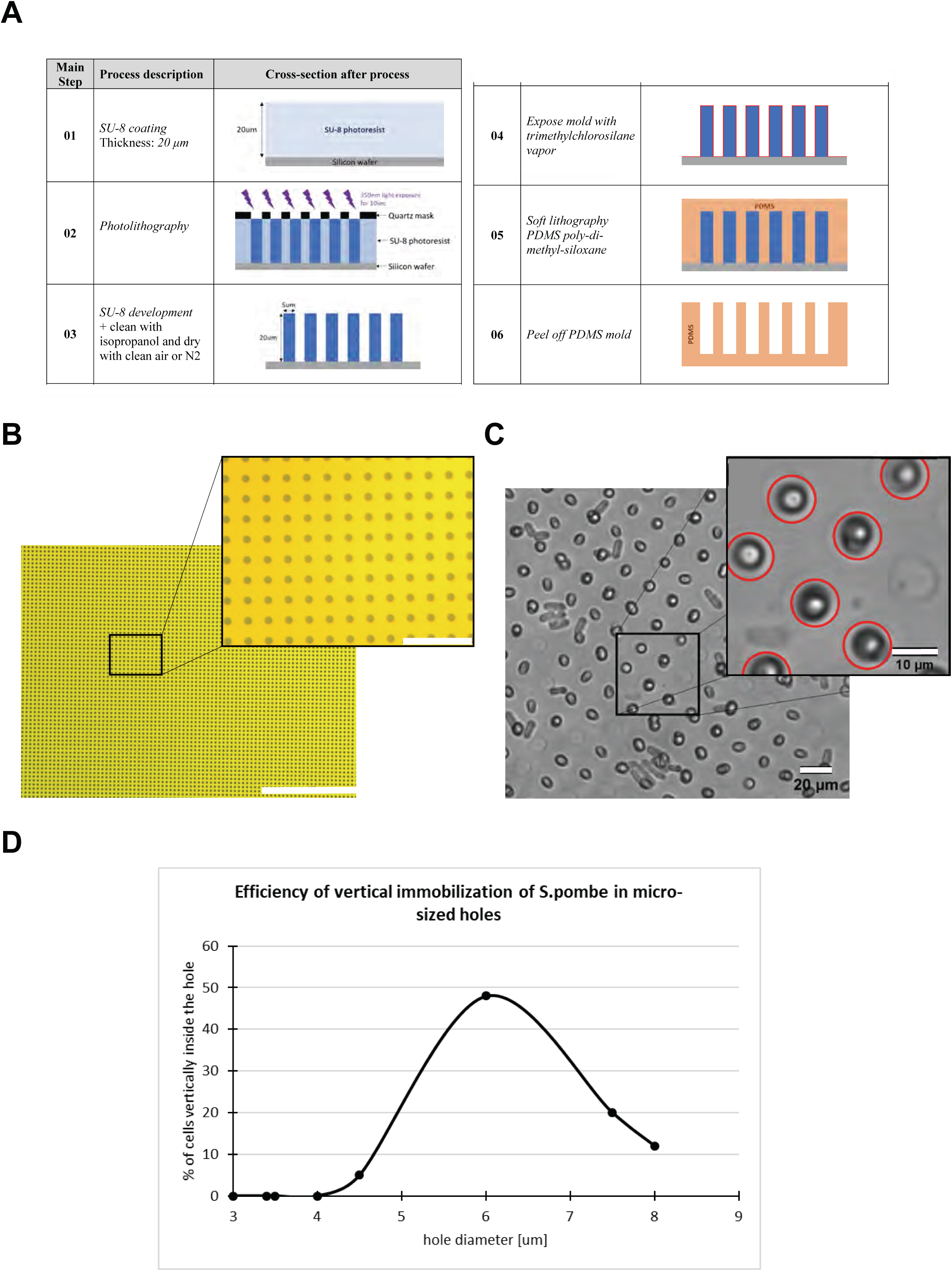
Fabrication of micro-holes for *S*. *pombe* cell vertical immobilization. **A.** Main steps of the process flow for the fabrication of the mold for the generation of the agarose support with micro-holes for vertical cell trapping. **B.** Wide-field image of the quartz mask for photolithography. Scale bar 400 µm and 40 µm (magnified view)**. C.** Visual inspection of the micro-hole provided the identification and count of cells correctly placed. Scale bar 20 µm and 10 µm (magnified view). **D.** To identify the optimal micro-hole diameter we fabricated different cell supports with the diameter size swept from 3 µm up to 8µm. We found that the diameter size at which the maximum number of cells were correctly verticalized was around 6µm.

**Figure 1 – supplement 2.**
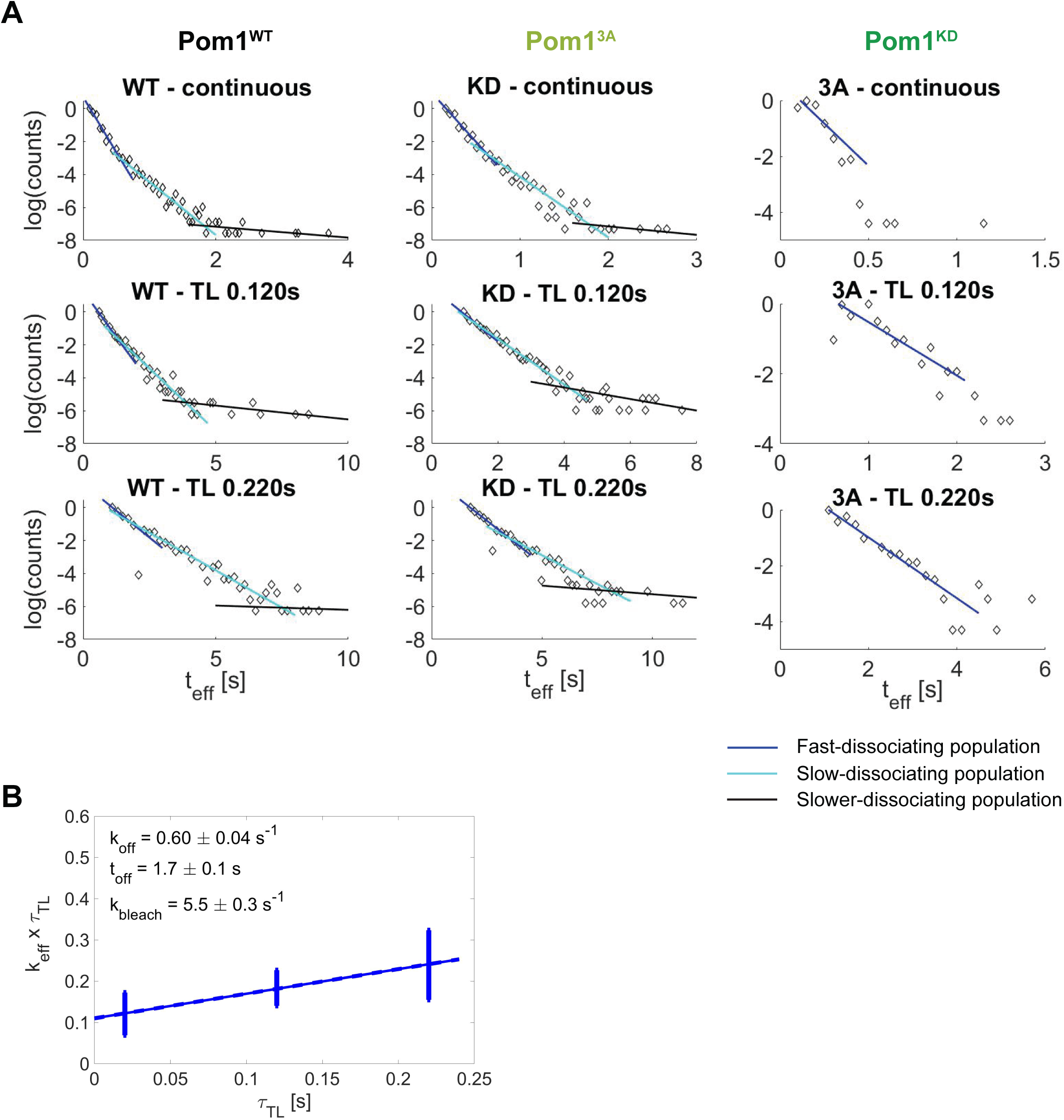
Pom1 dissociation analysis. **A.** Three-population identification and analysis through the fit of the exponential track length distribution. The last population (black) was excluded from the quantitative analysis due to the small sample size, but was used to limit the second population (cyan). **B.** Weighted linear fit of *k*_*eff*_ × τ_*TL*_ versus τ_*TL*_ for Pom1^3A^ from which the dissociation rate constant *k*_*off*_ and dissociation time constant *t*_*off*_ = 1/*k*_*off*_ are extracted according to Equation 2. Error bars correspond to the weights associated to each data point (S.D. from the fit of the exponential distribution of the track length).

**Figure 2 – supplement 1.**
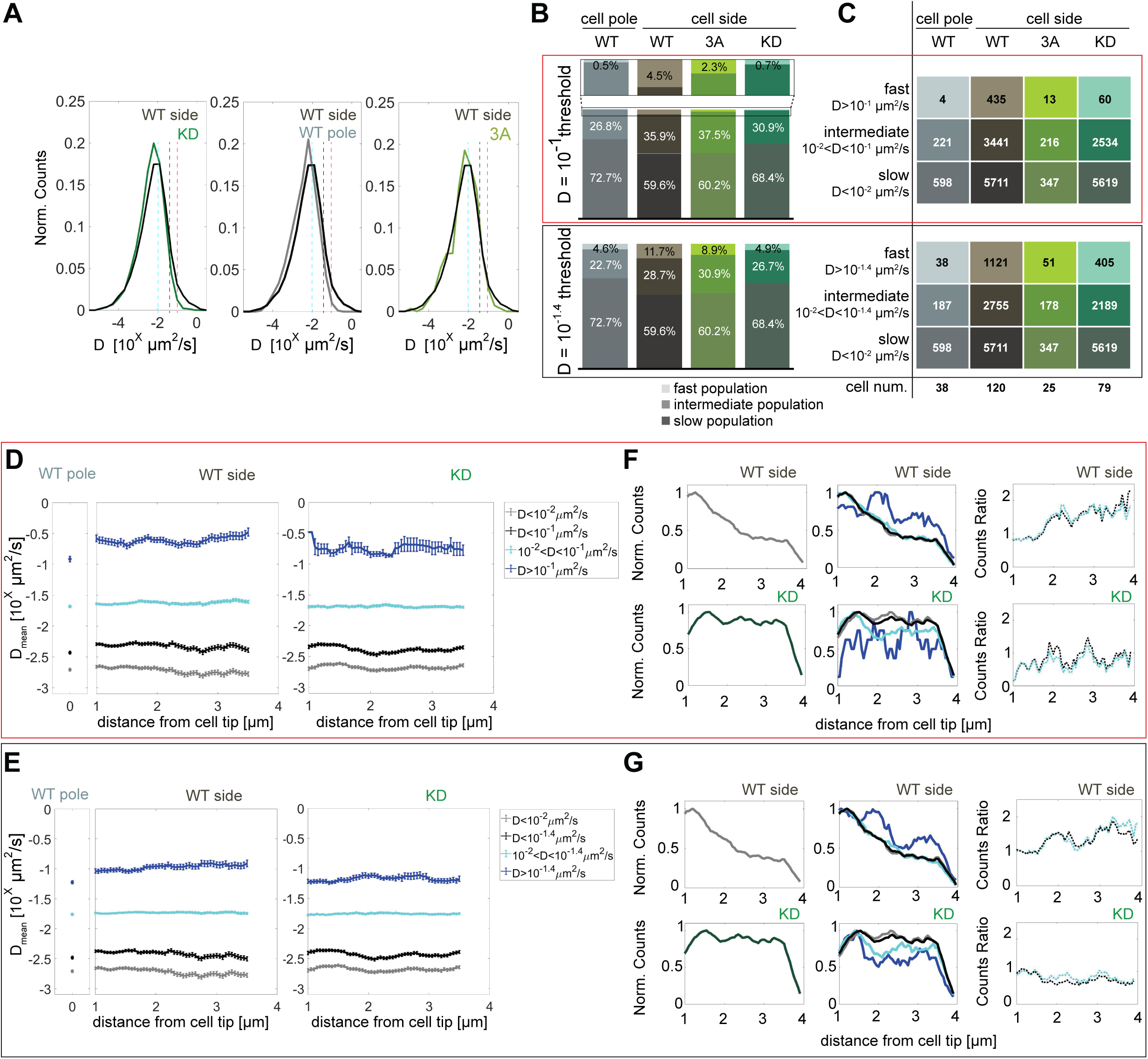
Pom1 diffusion coefficient analysis with different thresholds. **A.** Distribution of diffusion coefficients of all Pom1 molecules tracked in Pom1^WT^ (dark grey), Pom1^3A^ (light green) and Pom1^KD^ (green-cyan) at cell sides, and Pom1^WT^ at cell poles (light grey), as in Figure 2B. The two thresholds set to delimit the fast from the intermediate populations are represented by the vertical red (D = 10^−1^ µm^2^/s) and grey (D = 10^−1.4^ µm^2^/s) lines. The cyan line (D = 10^−2^ µm^2^/s) delimits the intermediate from the slow populations. **B.** Proportion of fast (light), intermediate and slow (dark) populations for Pom1^WT^, Pom1^3A^ and Pom1^KD^ at cell sides, and Pom1^WT^ at cell poles. **C.** Number of tracks and cells for each condition. **D-E.** Average diffusion coefficient as a function of the distance from the pole for fast (blue), intermediate (cyan), slow (grey) and intermediate plus slow (black) populations for Pom1^WT^ at the pole (left panel), Pom1^WT^ at the side (central panel) and Pom1^KD^ (right panel) strains. Error bar corresponds to standard error of the mean. **F-G.** Evolution of the total number of tracks for Pom1^WT^ (dark gray, top-left panel) and Pom1^KD^ (green-cyan, bottom-left panel) strains. Evolution of the number of tracks for fast (blue), intermediate (cyan), slow (grey) and slow plus intermediate (black) populations along the cell length for Pom1^WT^ (top-central panel) and Pom1^KD^ (bottom-central panel) strains. Ratios between the fast and intermediate population (cyan) and between fast and combined intermediate and slow population (black) for Pom1^WT^ (top-right panel) and Pom1^KD^ (bottom-right panel). For (B, C, D, F), the analysis in the red box uses D ≥ 10^−1^ µm^2^/s for the fast populations, 10^−2^ < D < 10^−1^ µm^2^/s for the intermediate populations and D < 10^−2^ µm^2^/s for the slow population. For (B, C, E, G) the analysis in the black box uses D ≥ 10^−1.4^ µm^2^/s for the fast populations, 10^−2^ < D < 10^−1.4^ µm^2^/s for the intermediate populations and D < 10^−2^ µm^2^/s for the slow population.

**Figure 2 – supplement 2.**
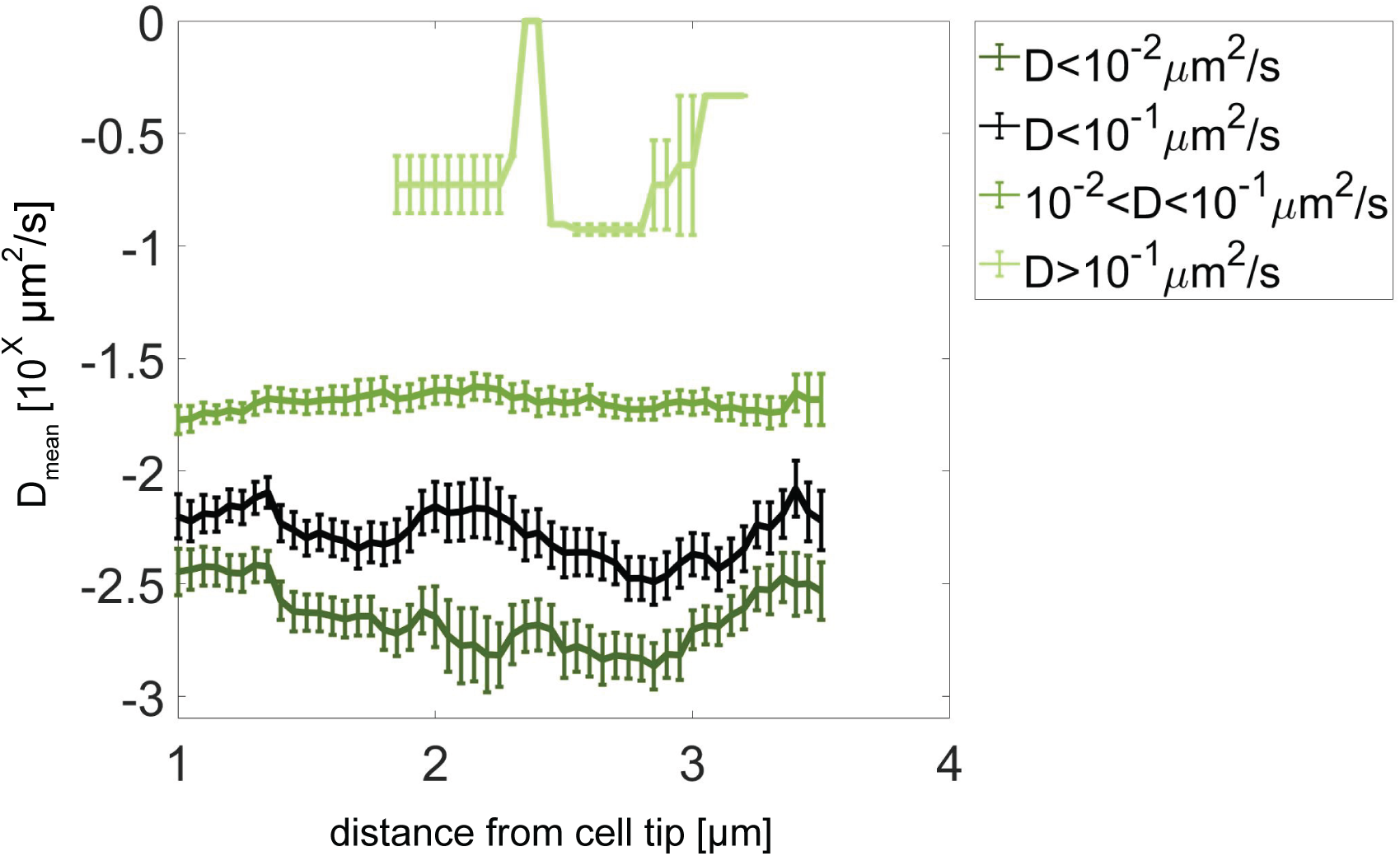
Pom1^3A^ diffusion coefficient analysis with different thresholds. Average diffusion coefficient as a function of the distance from the pole for fast population (light green, D ≥ 10^−1^ µm^2^/s), intermediate (medium green, 10^−2^ < D < 10^−1^ µm^2^/s), slow (dark green, D < 10^−2^ µm^2^/s), and slow plus intermediate (black, D < 10^−1^ µm^2^/s) populations. Error bar corresponds to standard error of the mean.

**Figure 3 – supplement 1:**
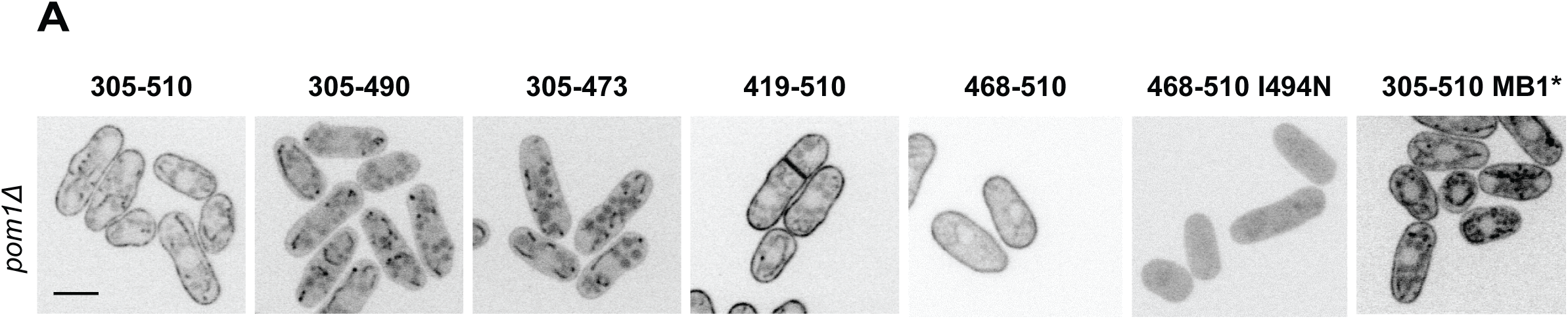
Analysis of Pom1 fragment localization in *pom1Δ* cells. Localization in *pom1Δ* cells of the Pom1 GFP-tagged fragments 1 to 8 presented in Figure 3A. Scale bar 5μm.

**Figure 4 – supplement 1:**
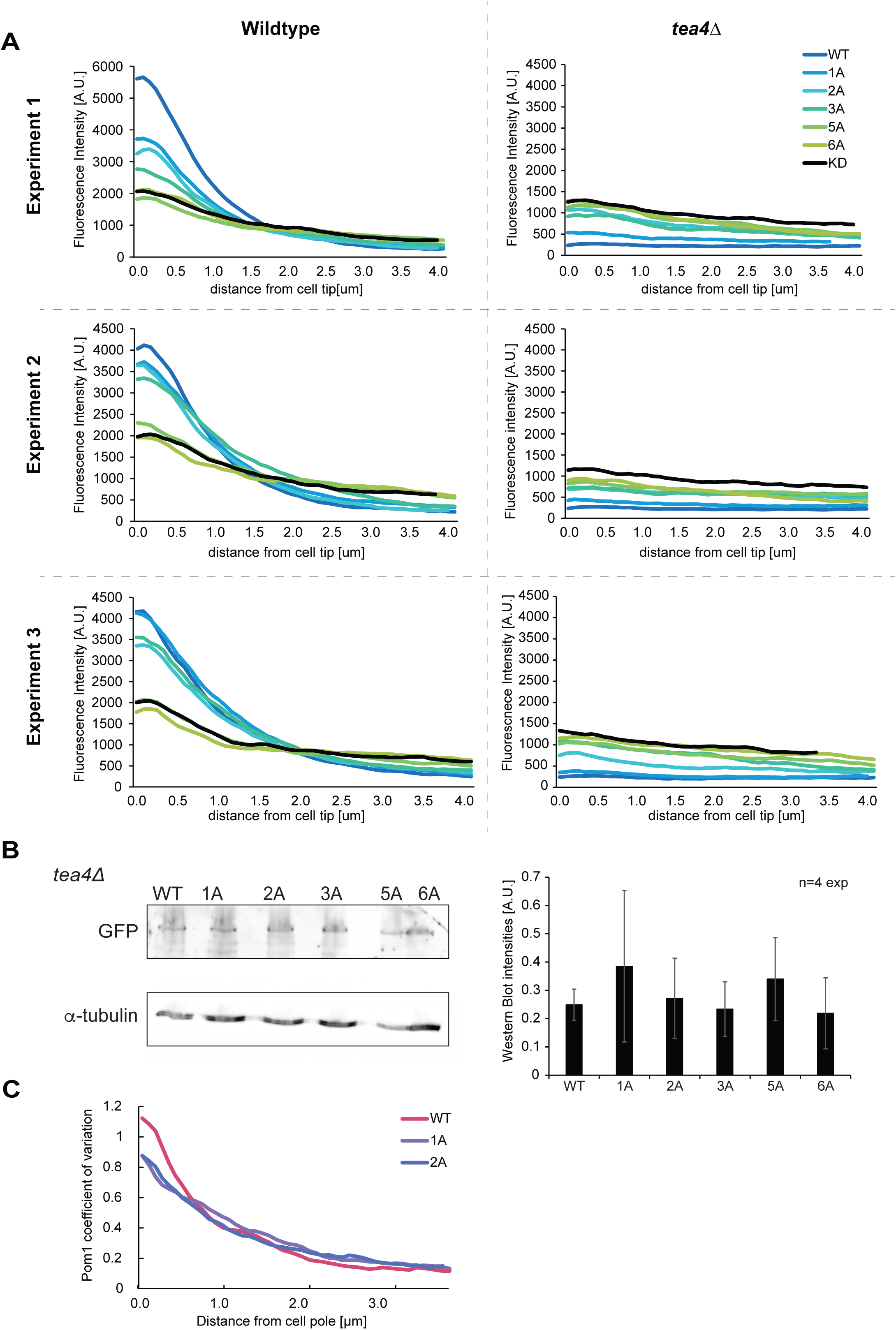
Analysis of Pom1 phospho-site mutant alleles. **A.** Individual experiments for the average fluorescence intensity plots of cortical gradient profiles, collected from time-average medial plane confocal images, shown in Figure 4B. Left: wild type background; right: *tea4Δ* cells. Graphs show averages of 80 gradient profiles per strain, collected in 20 individual cells per experiment. **B.** Western blot showing equal levels of expression of the various Pom1-GFP alleles in *tea4Δ* cells, as indicated. *α*-tubulin was used as a loading control. Average quantification of four experiments is shown on the right. Error bars: standard deviation. **C.** Coefficients of variation for Pom1^WT^, Pom1^1A^, and Pom1^2A^ from the gradient profiles shown in Figure 4B.

**Figure 5 – supplement 1:**
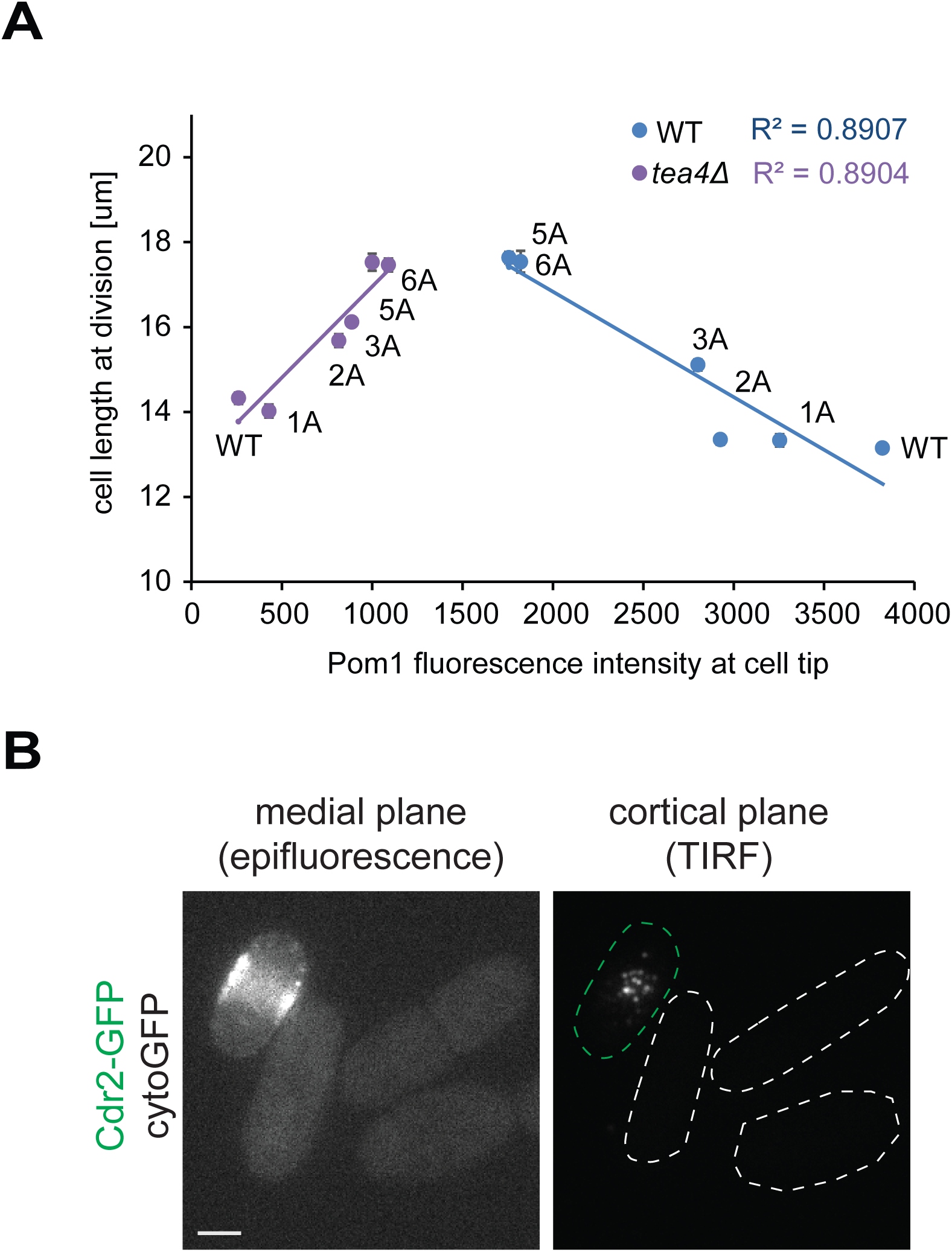
Pom1 levels at cell poles do not correlate with cell length at division. **A.** Correlation plot of cell length at division (values from Figure 5A) versus Pom1 intensity at cell poles (values from Figure 4B). Error bars are standard error. **B.** Mid-plane epifluorescence (left) and cortical TIRF (right) images of cytoGFP expressed from the *pom1* promoter. Note that the strain was mixed with a Cdr2-GFP strain (green dashed contour) to help find the TIRF focal plane. Scale bar 2.5μm.

**Figure 6 – supplement 1:**
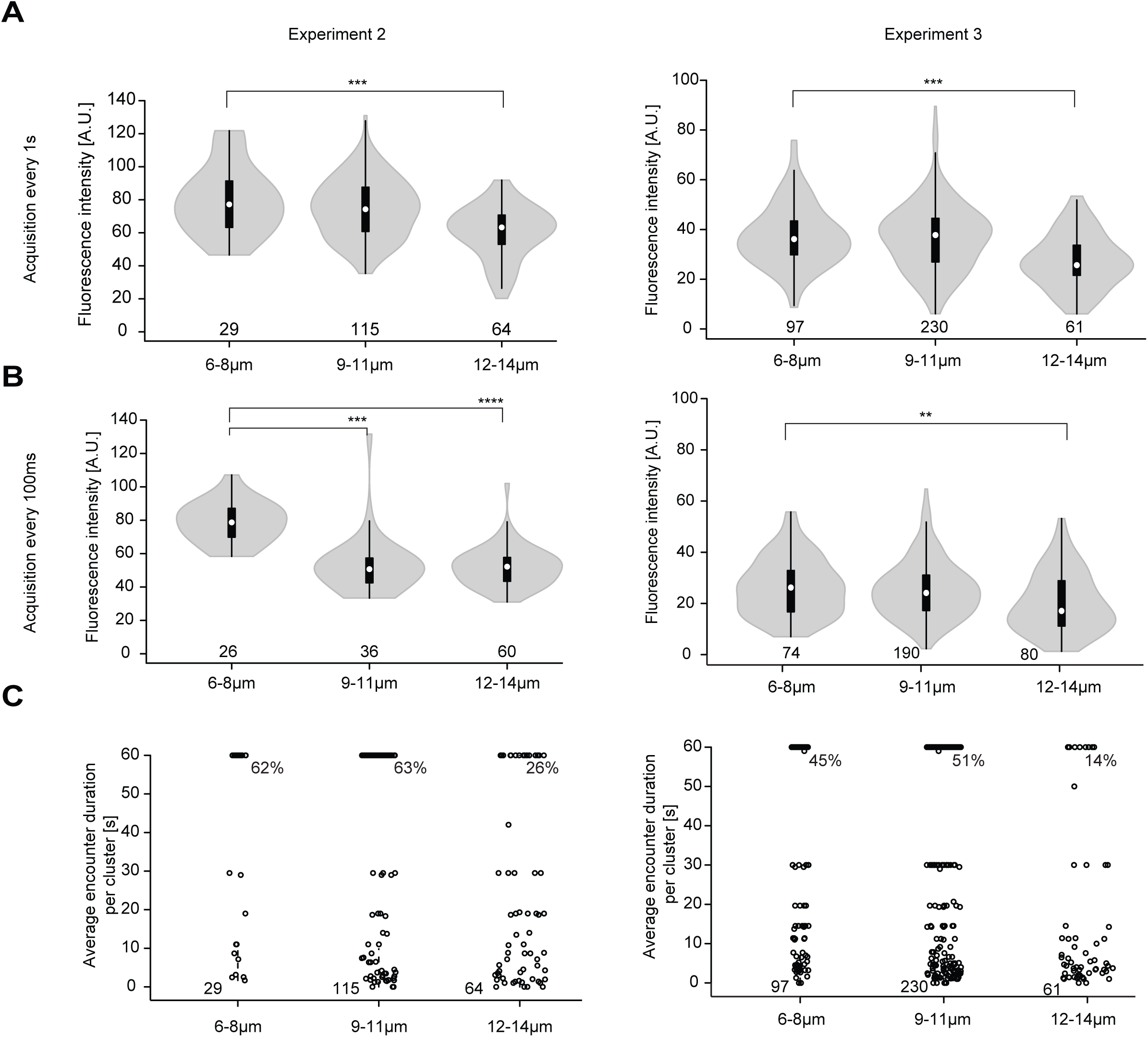
Replicate experiments showing higher Pom1-GFP fluorescence at Cdr2 nodes of short than long cells. A-B. Fluorescence intensity of Pom1-GFP at Cdr2 nodes measured as in Figure 6B every second over a 60s imaging period (A) or every 100ms over an 8s imaging period (B), sorted by cell length. **, p=10^−3^, ***, p≤10^−05^, ****, p≤10^−15^. **C.** Duration of individual Pom1 encounters with a Cdr2 node for cells of sorted length, measured as in Figure 6E. Note that 50 arbitrary units were chosen to define Pom1-GFP presence at a Cdr2 node. For experiment 2, the data was collected on the same day: for the 1 frame/s acquisition 29 clusters from 7 6-8μm long cells, 115 clusters from 15 9-11μm long cells, and 64 clusters from 8 12-14μm long cells were quantified; for the 100ms interval acquisition, 26 clusters from 6 6-8μm long cells, 36 clusters from 7 9-11μm long cells, and 60 clusters from 6 12-14μm long cells were quantified. For experiment 3, the data was collected on the same day: for the 1 frame/s acquisition 97 clusters from 16 6-8μm long cells, 230 clusters from 27 9-11μm long cells, and 62 clusters from 6 12-14μm long cells were quantified; for the 100ms interval acquisition 74 clusters from 14 6-8μm long cells, 190 clusters from 22 9-11μm long cells, and 80 clusters from 12 12-14μm long cells were quantified.

**Supplementary Video 1: Examples of Pom1-GFP cluster dynamics**. Pom1-GFP was imaged in TIRF mode every 100ms over a 8s imaging period. 4 clusters of interest are marked with a target with 3 prominent clusters visible at the start of the timelapse, and 1 smaller cluster at the bottom of the frame which appears at 0.2s and disappears temporarily at 1.2s and 1.8s and completely at 5.6s (possibly as it leaves the evanescent field on the side of the cell). The top positioned cluster splits at 2.8s (or even sooner), remerges at 3.3s, splits partially at 3.7s and completely at 4.4s, leaving behind 2 smaller clusters that move laterally and last until the end of the movie. The bottom cluster on at timepoint 0 is observed clearly until 6.9s. During this period, it dims and brightens, disperses and focalizes several times. At 6.9s disappears from the field of view, but may not be completely absent as a cluster (not marked) reappears at the same location at 7.0s. The cluster on the left moves laterally and splits in two clusters at 2.2s and re-merges at 2.7s and remains trackable until the end of the 8s imaging period. All clusters tracked in this movie show examples of dimming, followed by brightening of the signal. The movie is shown in quasi-real time.

**Supplementary Video 2: Examples of Pom1-GFP cluster dynamics.** Pom1-GFP was imaged in TIRF mode every 100ms over a 8s imaging period. 3 clusters of interest are marked with a target. The cluster on the left-splits in 3 smaller clusters at 0.1s, after which the clusters appear to detach. The cluster on the top right splits in 2 clusters at 0.1s and re-merges briefly at 0.2s, after which the cluster increases in brightness and splits again at 0.6s. The small cluster on the left moves laterally, dims in fluorescence and eventually detaches at 1.3s. The right top cluster and the bottom cluster, marked from the beginning of the movie move laterally, occasionally splitting and merging. The top cluster splits at 2.4s and re-merges at 2.7s, as does the bottom cluster at 3.2s and 3.4s, and 4.5s. Eventually, the clusters will merge at 4.9s, forming a bright and stable cluster, which remains trackable until the end of the 8s imaging period. The movie is shown in quasi-real time.

## Supplementary Notes

### Data analysis

The emerging of deviation from a linear trend in the log plot of the distribution of the track lengths indicated the possible presence of transient binding with different kinetic constants. Assuming the presence of at least two populations, one with a faster off-rate constant and one with a slower one, the model describing the dissociation of Pom1 then becomes:

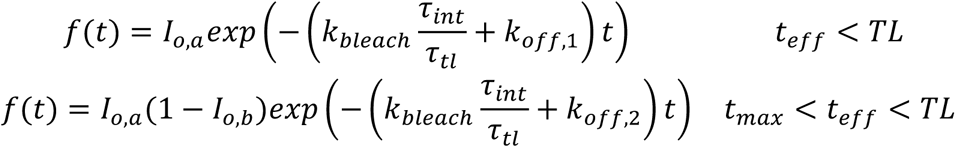

Where *I*_*o,a*_ is the number of molecules at start, *t* is real time in seconds and *I*_*o,b*_ represents the percentage of Pom1 molecules exhibiting off-rate constant *k*_off,2_.

### Acquisition setup

#### Fluorescence channel

- Camera: EMCCD Prime 95b
- readout rate = 10MHz 16-bits
- Exposure time: 20ms
- Gain: 1000
- Focus locked by CRISP-Focus
- Excitation 560nm : “AOTF-DAC1-Wavelength nm” = 30mW
- “AOTF-DAC1-Volts”: “3”
- Activation 405nm: “AOTF-DAC3-Wavelength nm”
- “AOTF-DAC3-Volts” = 0 to 5

#### Phase-contrast channel

- Camera: TIS-DCAM
- Exposure time: 50ms

---

**Table S1.** *S*.*pombe* strains used in this study.

| Number | Genotype | Source |
| --- | --- | --- |
| **Figure 1-2** |  |  |
| YSM3262 | h- pom1-mEos3.2-kanMX ade6-M216 leu1-32 ura4-D18 | This study. |
| YSM3263 | h+ pom1<sup>KD</sup>-mEos3.2-kanMX leu1-32 ura4-D18 | This study. |
| YSM3264 | h+ pom1<sup>3A(2,4,5)</sup>-mEos3.2-kanMX ade6-M216 leu1-32 ura4-D18 | This study. |
| **Figure 3** |  |  |
| YSM3265 | h- ura4-D18::ppom1-pom1<sub>(305-510)</sub>-GFP::ura4+ ade6-M210 leu1-32 | This study. |
| YSM3266 | h- ura4-D18::ppom1-pom1<sub>(305-490)</sub>-GFP::ura4+ ade6-M210 leu1-32 | This study. |
| YSM3267 | h- ura4-D18::ppom1-pom1<sub>(305-473)</sub>-GFP::ura4+ ade6-M210 leu1-32 | This study. |
| YSM3268 | h- ura4-D18::ppom1-pom1<sub>(419-510)</sub>-GFP::ura4+ ade6-M210 leu1-32 | This study. |
| YSM3269 | h- ura4-D18::ppom1-pom1<sub>(468-510)</sub>-GFP::ura4+ ade6-M210 leu1-32 | This study. |
| YSM3270 | h- ura4-D18::ppom1-pom1<sub>(468-510)</sub><sup>I494N</sup>-GFP::ura4+ ade6-M210 leu1-32 | This study. |
| YSM3271 | h- ura4-D18::ppom1-pom1<sub>(305-510)</sub><sup>I494N</sup>-GFP::ura4+ ade6-M210 leu1-32 | This study. |
| YSM3272 | h- ura4-D18::ppom1-pom1<sub>(305-510)</sub><sup>7ALA(MB1*)</sup>-GFP::ura4+ ade6-M210 leu1-32 | This study. |
| YSM3273 | h- ura4-D18::ppom1-pom1<sub>(305-510)</sub><sup>I494N-MB1*</sup>-GFP::ura4+ ade6-M210 leu1-32 | This study. |
| YSM3274 | h- pom1-eGFP-kanMX ade6-M210 leu1-32 ura4-D18 | This study. |
| YSM3275 | h- pom1<sup>I494N</sup>-eGFP-kanMX ade6-M210 leu1-32 ura4-D18 | This study. |
| YSM3276 | h- pom1<sup>7ALA(MB1*)</sup>-eGFP-kanMX ade6-M210 leu1-32 ura4-D18 | This study. |
| YSM3277 | h- pom1<sup>I494N-MB1*</sup>-eGFP-kanMX ade6-M210 leu1-32 ura4-D18 | This study. |
| YSM3278 | h- pom1<sup>I494N-MB1*-5PxxP*</sup>-eGFP-kanMX ade6-M210 leu1-32 ura4-D18 | This study. |
| **Figure S3** |  |  |
| YSM3279 | pom1Δ::kanMX ura4-D18::pom1<sub>(305-510)</sub>-GFP::ura4+ ade6-M210 leu1-32 | This study. |
| YSM3280 | pom1Δ::kanMX ura4-D18::pom1<sub>(305-490)</sub>-GFP::ura4+ ade6-M210 leu1-32 | This study. |
| YSM3281 | pom1Δ::kanMX ura4-D18::pom1<sub>(305-473)</sub>-GFP::ura4+ ade6-M210 leu1-32 | This study. |
| YSM3282 | pom1Δ::kanMX ura4-D18::pom1<sub>(419-510)</sub>-GFP::ura4+ ade6-M210 leu1-32 | This study. |
| YSM3283 | pom1Δ::kanMX ura4-D18::pom1<sub>(468-510)</sub>-GFP::ura4+ ade6-M210 leu1-32 | This study. |
| YSM3284 | pom1Δ::kanMX ura4-D18::pom1<sub>(468-510)</sub><sup>I494N</sup>-GFP::ura4+ ade6-M210 leu1-32 | This study. |
| YSM3285 | pom1Δ::kanMX ura4-D18::pom1<sub>(305-510)</sub><sup>7ALA(MB1*)</sup>-GFP::ura4+ ade6-M210 leu1-32 | This study. |
| **Figure 4** |  |  |
| YSM1912 | h- pom1-GFP-kanMX ade6-M216 leu1-32 ura4-D18 | (Hachet et al., 2011) |
| YSM3287 | h+ pom1<sup>1A(5)</sup>-GFP-kanMX ade6-M216 leu1-32 ura4-D18 | This study. |
| YSM3288 | h+ pom1<sup>2A(4,5)</sup>-GFP-kanMX ade6-M216 leu1-32 ura4-D18 | This study. |
| YSM3289 | h+ pom1<sup>3A(2,4,5)</sup>-GFP-kanMX ade6-M216 leu1-32 ura4-D18 | This study. |
| YSM3290 | h+ pom1<sup>5A(1,2,3,4,5)</sup>-GFP-kanMX ade6-M216 leu1-32 ura4-D18 | This study. |
| YSM2271 | h+ pom1<sup>6A(1,2,3,4,5,6)</sup>-GFP-kanMX ade6-M216 leu1-32 ura4-D18 | (Hachet et al., 2011) |
| YSM1329 | h- pom1<sup>KD</sup>-GFP-kanMX ade6-M216 leu1-32 ura4-D18 | (Hachet et al., 2011) |
| YSM3291 | tea4Δ::hphMX pom1-GFP-kanMX ade6-M216 leu1-32 ura4-D18 | This study. |
| YSM3292 | tea4Δ::hphMX pom1<sup>1A(5)</sup>-GFP-kanMX ade6-M216 leu1-32 ura4-D18 | This study. |
| YSM3293 | tea4Δ::hphMX pom1<sup>2A(4,5)</sup>-GFP-kanMX ade6-M216 leu1-32 ura4-D18 | This study. |
| YSM3294 | tea4Δ::hphMX pom1<sup>3A(2,4,5)</sup>-GFP-kanMX ade6-M216 leu1-32 ura4-D18 | This study. |
| YSM3295 | tea4Δ::hphMX pom1<sup>5A(1,2,3,4,5)</sup>-GFP-kanMX ade6-M216 leu1-32 ura4-D18 | This study. |
| YSM3296 | tea4Δ::hphMX pom1<sup>6A(1,2,3,4,5,6)</sup>-GFP-kanMX ade6-M216 leu1-32 ura4-D18 | This study. |
| YSM1855 | tea4Δ::kanMX pom1<sup>KD</sup>-GFP-kanMX ade6-M210 leu1-32 ura4-D18 | (Hachet et al., 2011) |
| YSM3274 | h- pom1-eGFP-kanMX ade6-M210 leu1-32 ura4-D18 | This study. |
| YSM3276 | h- pom1<sup>7ALA(MB1*)</sup>-eGFP-kanMX ade6-M210 leu1-32 ura4-D18 | This study. |
| Ysm3286 | h- pom1<sup>7ALA(MB1*)-1(A5)</sup>-eGFP-kanMX ade6-M210 leu1-32 ura4-D18 | This study. |
| **Figure 5** |  |  |
| YSM1912 | h- pom1-GFP-kanMX ade6-M216 leu1-32 ura4-D18 | (Hachet et al., 2011) |
| YSM3289 | h+ pom1<sup>3A(2,4,5)</sup>-GFP-kanMX ade6-M216 leu1-32 ura4-D18 | This study. |
| YSM1413 | h- cdr2-GFP-ura4+ ade6-M216 leu1-32 ura4-D18 | (Bhatia et al., 2013) |
| YMM773 | h- ura4-D18::ppom1-sfGFP-tCYC1::ura4+ ade+ leu+ | This study. |
| YSM165 | h- tea4Δ::kanMX pom1-GFP-kanMX ade6-M210 leu1-32 ura4-D18 | (Martin et al., 2005) |
| **Figure 6** |  |  |
| YSM1457 | pom1-GFP-kanMX cdr2-tdTomato-natMX | This study. |

**Table S2.** Plasmid list.

| Number | Description | Purpose |
| --- | --- | --- |
| pSM2114 | pAV0133-*ppom1*-pom1<sub>(305-510)</sub>-GFP | *ura4* integration |
| pSM2115 | pAV0133-*ppom1*-pom1<sub>(305-490)</sub>-GFP | *ura4* integration |
| pSM2116 | pAV0133-*ppom1*-pom1<sub>(305-510)</sub><sup>7ALA(MB1*)</sup>-GFP | *ura4* integration |
| pSM2117 | pAV0133-*ppom1*-pom1<sub>(305-473)</sub>-GFP | *ura4* integration |
| pSM2118 | pAV0133-*ppom1*-pom1<sub>(419-510)</sub>-GFP | *ura4* integration |
| pSM1848 | pAV0133-*ppom1*-pom1<sub>(419-510)</sub>-GFP | *ura4* integration |
| pSM1850 | pAV0133-*ppom1*-pom1<sub>(419-510)</sub><sup>I494N</sup>-GFP | *ura4* integration |
| pSM2132 | pAV0133-*ppom1*-pom1<sub>(305-510)</sub><sup>I494N</sup>-GFP | *ura4* integration |
| pSM2133 | pAV0133-*ppom1*-pom1<sub>(305-510)</sub><sup>MB1*-I494N</sup>-GFP | *ura4* integration |
| pSM2142 | pFA6A-*ppom1*-pom1ORF-eGFP-kanMX- *pom1* 3'UTR | target endogenous locus |
| pSM2146 | pFA6A-*ppom1*-pom1<sup>I494N</sup>-eGFP-kanMX- *pom1* 3'UTR | target endogenous locus |
| pSM2237 | pFA6A-*ppom1*-pom1<sup>MB1*</sup>-eGFP-kanMX- *pom1* 3'UTR | target endogenous locus |
| pSM2238 | pFA6A-*ppom1*-pom1<sup>MB1*-I494N</sup>-eGFP-kanMX- *pom1* 3'UTR | target endogenous locus |
| pSM2264 | pFA6A-*ppom1*-pom1<sup>MB1*-1A(5)</sup>-eGFP-kanMX- *pom1* 3'UTR | target endogenous locus |
| pSM2328 | pFA6A-*ppom1*-pom1<sup>MB1*-I494N-5PxxP*</sup>-eGFP-kanMX-3'UTR | target endogenous locus |
| pSM738 | pREP41-pom1-GFP | phosphosite mutagenesis |
| pSM1502 | pREP41-pom1<sup>1A(5)</sup> | phosphosite mutagenesis |
| pSM1576 | pREP41-pom1<sup>2A(2,5)</sup> | phosphosite mutagenesis |
| pSM1527 | pREP41-pom1<sup>3A(2,4,5)</sup> | phosphosite mutagenesis |
| pSM1866 | pREP41-pom1<sup>5A(1,2,3,4,5)</sup> | phosphosite mutagenesis |
| pSM1731 | pFA6A- pom1<sub>(911-1087)</sub>-GFP-kanMX-*pom1* 3'UTR | C-terminal GFP tagging |

**Table S3.**
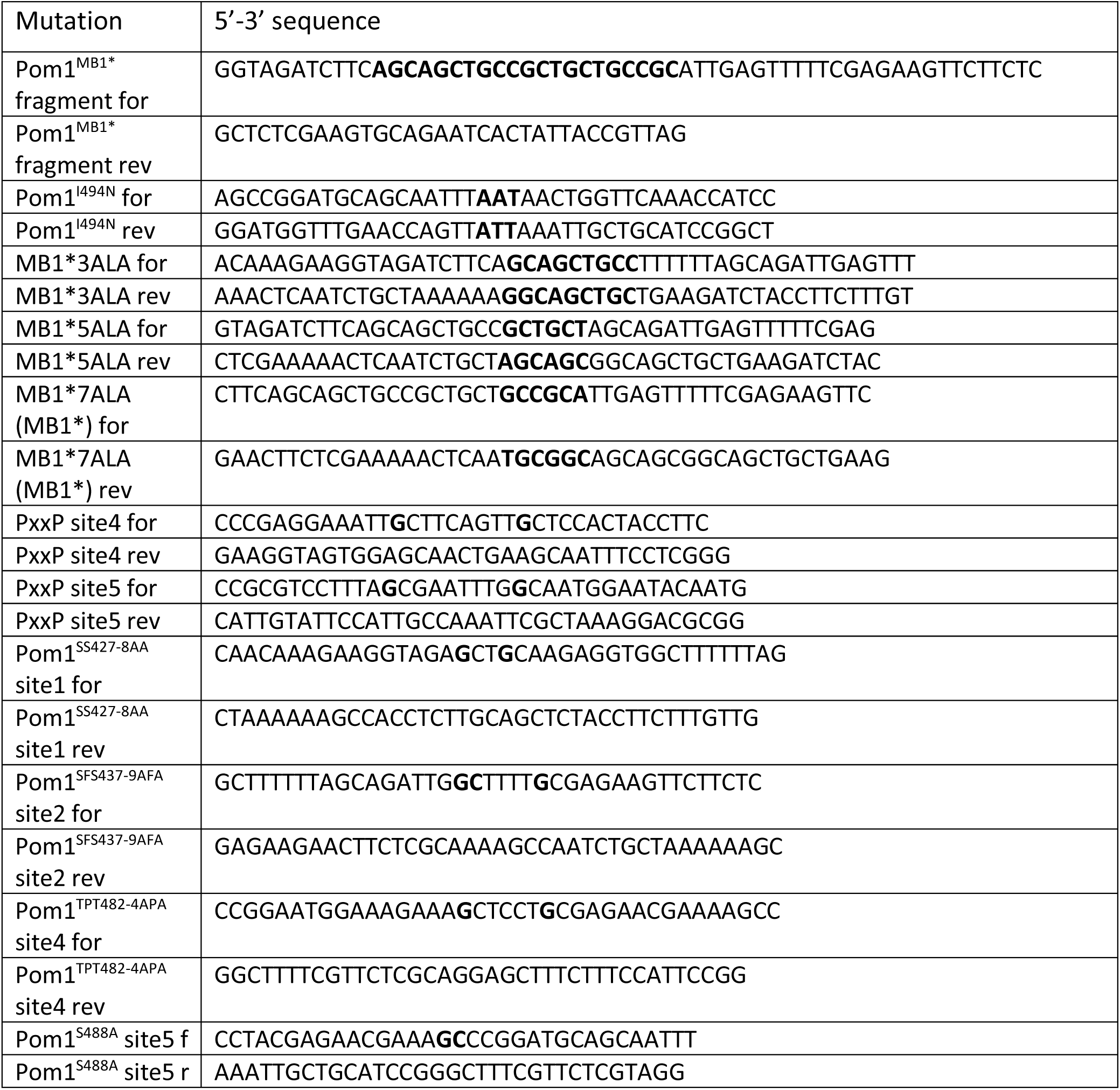
Primers used for mutagenesis. Bold residues indicate changes from the wildtype sequence.

**Table S4.**
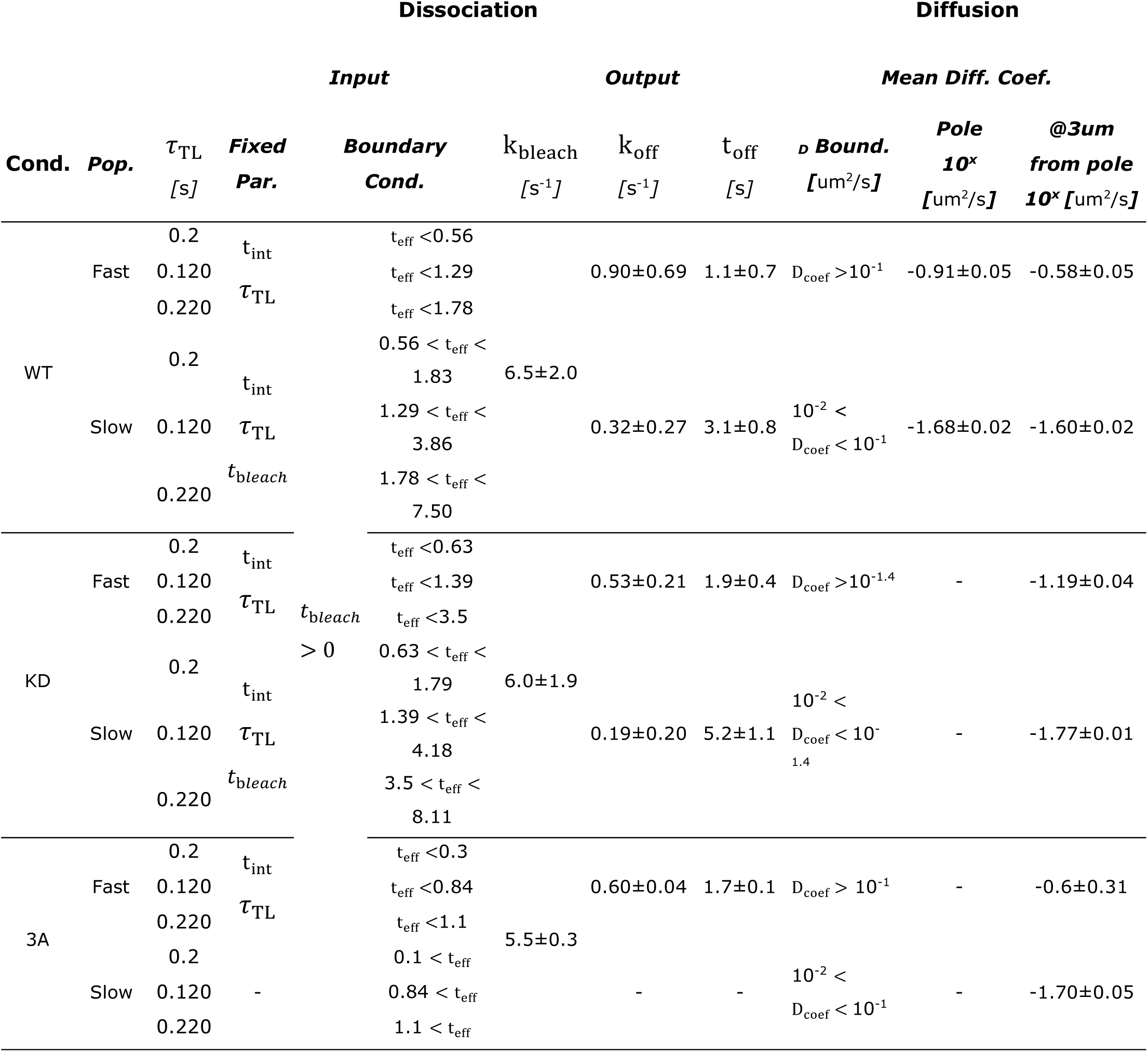
Global fitting parameters and outputs.

## References

Allard, C.A.H., H.E. Opalko, K.W. Liu, U. Medoh, and J.B. Moseley. 2018. Cell size-dependent regulation of Wee1 localization by Cdr2 cortical nodes. J Cell Biol. 217:1589–1599.

Alvarez-Tabares, I., A. Grallert, J.M. Ortiz, and I.M. Hagan. 2007. Schizosaccharomyces pombe protein phosphatase 1 in mitosis, endocytosis and a partnership with Wsh3/Tea4 to control polarised growth. J Cell Sci. 120:3589–3601.

Bähler, J., and P. Nurse. 2001. Fission yeast Pom1p kinase activity is cell cycle regulated and essential for cellular symmetry during growth and division. Embo J. 20:1064–1073.

Bähler, J., and J.R. Pringle. 1998. Pom1p, a fission yeast protein kinase that provides positional information for both polarized growth and cytokinesis. Genes Dev. 12:1356–1370.

Bähler, J., J.Q. Wu, M.S. Longtine, N.G. Shah, A. McKenzie, 3rd, A.B. Steever, A. Wach, P. Philippsen, and J.R. Pringle. 1998. Heterologous modules for efficient and versatile PCR-based gene targeting in Schizosaccharomyces pombe. Yeast. 14:943–951.

Bhatia, P., O. Hachet, M. Hersch, S.A. Rincon, M. Berthelot-Grosjean, S. Dalessi, L. Basterra, S. Bergmann, A. Paoletti, and S.G. Martin. 2013. Distinct levels in Pom1 gradients limit Cdr2 activity and localization to time and position division. Cell Cycle. 13.

Briscoe, J., and S. Small. 2015. Morphogen rules: design principles of gradient-mediated embryo patterning. Development. 142:3996–4009.

Brzeska, H., J. Guag, K. Remmert, S. Chacko, and E.D. Korn. 2010. An experimentally based computer search identifies unstructured membrane-binding sites in proteins: application to class I myosins, PAKS, and CARMIL. J Biol Chem. 285:5738–5747.

Carlini, L., S.J. Holden, K.M. Douglass, and S. Manley. 2015. Correction of a Depth-Dependent Lateral Distortion in 3D Super-Resolution Imaging. PLoS One. 10:e0142949.

Crocker, J.C., and D.G. Grier. 1996a. Methods of Digital Video Microscopy for Colloidal Studies. Journal of Colloidal and Interface Science. 179:298–310.

Crocker, J.C., and D.G. Grier. 1996b. When Like Charges Attract: The Effects of Geometrical Confinement on Long-Range Colloidal Interactions. Physical review letters. 77:1897–1900.

Deng, L., S. Baldissard, A.N. Kettenbach, S.A. Gerber, and J.B. Moseley. 2014. Dueling kinases regulate cell size at division through the SAD kinase Cdr2. Curr Biol. 24:428–433.

Facchetti G, Knapp B, Flor-Parra I, Chang F, and H. M. 2019. Reprogramming Cdr2-Dependent Geometry-Based Cell Size Control in Fission Yeast. Curr Biol.

Gebhardt, J.C., D.M. Suter, R. Roy, Z.W. Zhao, A.R. Chapman, S. Basu, T. Maniatis, and X.S. Xie. 2013. Single-molecule imaging of transcription factor binding to DNA in live mammalian cells. Nat Methods. 10:421–426.

Gerganova, V., and S.G. Martin. 2018. Dynamic visits of cortical structures probe for cell size. J Cell Biol. 217:1559–1561.

Hachet, O., M. Berthelot-Grosjean, K. Kokkoris, V. Vincenzetti, J. Moosbrugger, and S.G. Martin. 2011. A phosphorylation cycle shapes gradients of the DYRK family kinase Pom1 at the plasma membrane. Cell. 145:1116–1128.

Hersch, M., O. Hachet, S. Dalessi, P. Ullal, P. Bhatia, S. Bergmann, and S.G. Martin. 2015. Pom1 gradient buffering through intermolecular auto-phosphorylation. Molecular systems biology. 11:818.

Holden, S.J., T. Pengo, K.L. Meibom, C. Fernandez Fernandez, J. Collier, and S. Manley. 2014. High throughput 3D super-resolution microscopy reveals Caulobacter crescentus in vivo Z-ring organization. Proc Natl Acad Sci U S A. 111:4566–4571.

Huang, B., W. Wang, M. Bates, and X. Zhuang. 2008. Three-dimensional super-resolution imaging by stochastic optical reconstruction microscopy. Science. 319:810–813.

Kanoh, J., and P. Russell. 1998. The protein kinase Cdr2, related to Nim1/Cdr1 mitotic inducer, regulates the onset of mitosis in fission yeast. Mol Biol Cell. 9:3321–3334.

Kelkar, M., and S.G. Martin. 2015. PKA antagonizes CLASP-dependent microtubule stabilization to relocalize Pom1 and buffer cell size upon glucose limitation. Nat Commun. 6:8445.

Kokkoris, K., D.G. Castro, and S.G. Martin. 2014. Tea4-phosphatase I landmark promotes local growth by dual Cdc42 GEF recruitment and GAP exclusion. J Cell Sci.

Kretschmer, S., and P. Schwille. 2016. Pattern formation on membranes and its role in bacterial cell division. Curr Opin Cell Biol. 38:52–59.

Laplante, C., F. Huang, J. Bewersdorf, and T.D. Pollard. 2016. High-Speed Super-Resolution Imaging of Live Fission Yeast Cells. Methods Mol Biol. 1369:45–57.

Manley, S., J.M. Gillette, G.H. Patterson, H. Shroff, H.F. Hess, E. Betzig, and J. Lippincott-Schwartz. 2008. High-density mapping of single-molecule trajectories with photoactivated localization microscopy. Nat Methods. 5:155–157.

Martin, S.G., and M. Berthelot-Grosjean. 2009. Polar gradients of the DYRK-family kinase Pom1 couple cell length with the cell cycle. Nature. 459:852–856.

Martin, S.G., W.H. McDonald, J.R. Yates, 3rd, and F. Chang. 2005. Tea4p links microtubule plus ends with the formin for3p in the establishment of cell polarity. Dev Cell. 8:479–491.

Morrell, J.L., C.B. Nichols, and K.L. Gould. 2004. The GIN4 family kinase, Cdr2p, acts independently of septins in fission yeast. J Cell Sci. 117:5293–5302.

Moseley, J.B., A. Mayeux, A. Paoletti, and P. Nurse. 2009. A spatial gradient coordinates cell size and mitotic entry in fission yeast. Nature. 459:857–860.

Padte, N.N., S.G. Martin, M. Howard, and F. Chang. 2006. The cell-end factor pom1p inhibits mid1p in specification of the cell division plane in fission yeast. Curr Biol. 16:2480–2487.

Pan, K.Z., T.E. Saunders, I. Flor-Parra, M. Howard, and F. Chang. 2014. Cortical regulation of cell size by a sizer cdr2p. eLife. 3:e02040.

Rincon, S.A., P. Bhatia, C. Bicho, M. Guzman-Vendrell, V. Fraisier, W.E. Borek, F.D. Alves, F. Dingli, D. Loew, J. Rappsilber, K.E. Sawin, S.G. Martin, and A. Paoletti. 2014. Pom1 regulates the assembly of Cdr2-Mid1 cortical nodes for robust spatial control of cytokinesis. J Cell Biol.

Saunders, T.E., K.Z. Pan, A. Angel, Y. Guan, J.V. Shah, M. Howard, and F. Chang. 2012. Noise reduction in the intracellular pom1p gradient by a dynamic clustering mechanism. Dev Cell. 22:558–572.

Tarantino, N., J.Y. Tinevez, E.F. Crowell, B. Boisson, R. Henriques, M. Mhlanga, F. Agou, A. Israel, and E. Laplantine. 2014. TNF and IL-1 exhibit distinct ubiquitin requirements for inducing NEMO-IKK supramolecular structures. J Cell Biol. 204:231–245.

Tatebe, H., K. Shimada, S. Uzawa, S. Morigasaki, and K. Shiozaki. 2005. Wsh3/Tea4 is a novel cell-end factor essential for bipolar distribution of Tea1 and protects cell polarity under environmental stress in S. pombe. Curr Biol. 15:1006–1015.

Wang, L., and P.T. Tran. 2014. Visualizing single rod-shaped fission yeast vertically in micro-sized holes on agarose pad made by soft lithography. Methods Cell Biol. 120:227–234.

Wolter, S., A. Loschberger, T. Holm, S. Aufmkolk, M.C. Dabauvalle, S. van de Linde, and M. Sauer. 2012. rapidSTORM: accurate, fast open-source software for localization microscopy. Nat Methods. 9:1040–1041.

Wood, E., and P. Nurse. 2013. Pom1 and cell size homeostasis in fission yeast. Cell Cycle. 12.

Young, P.G., and P.A. Fantes. 1987. Schizosaccharomyces pombe mutants affected in their division response to starvation. J Cell Sci. 88 (Pt 3):295–304.

